# mir124-dependent tagging of glutamatergic synapses by synaptopodin controls non-uniform and input-specific homeostatic synaptic plasticity

**DOI:** 10.1101/2021.04.30.442089

**Authors:** Sandra Dubes, Anaïs Soula, Sébastien Benquet, Béatrice Tessier, Christel Poujol, Alexandre Favereaux, Olivier Thoumine, Mathieu Letellier

## Abstract

Homeostatic synaptic plasticity (HSP) is a process by which neurons adjust synaptic strengths to compensate for various perturbations and which allows to stabilize neuronal activity. Yet, whether the highly diverse synapses harboring a neuron respond uniformly to a same perturbation is unclear and the underlying molecular determinants remain to be identified. Here, using patch-clamp recordings, immunolabeling and imaging approaches, we report that the ability of individual synapses to undergo HSP in response to activity-deprivation paradigms depends on the local expression of the spine apparatus related protein synaptopodin (SP) acting as a synaptic tag to promote AMPA receptor synaptic accumulation and spine growth. Gain and loss-of-function experiments indicate that this process relies on the local de-repression of SP translation by miR124 which supports both non-uniform and synapse-autonomous HSP induced by global or input-specific activity deprivation, respectively. Our findings uncover an unexpected synaptic-tagging mechanism for HSP, whose molecular actors are intriguingly shared with Hebbian plasticity and linked to multiple neurological diseases.

## Introduction

In the face of continuous alterations of neuronal activity, neurons adjust the efficacy of their connections in order to maintain stable network activity, a process referred to as homeostatic synaptic plasticity (HSP). These homeostatic adjustments are achieved at least in part through changes in postsynaptic AMPA receptor (AMPAR) number and function. One major form of HSP is multiplicative synaptic scaling, in which the synaptic strength of every excitatory synapses on a neuron is slowly scaled up or down with a same gain to compensate for prolonged alterations of neuronal firing rate (Turrigiano, 2008). In classical experimental paradigms, chronic treatment of primary neurons with tetrodotoxin (TTX) or bicuculline to inhibit or enhance neuronal activity, respectively, induces a uniform and widespread increase or decrease of postsynaptic strengths (Gainey et al., 2009; Turrigiano et al., 1998). This adaptation involves the synthesis of proteins as diverse as glutamatergic receptors, scaffolding proteins, voltage-gated ion channels, kinases, secreted factors and cell adhesion molecules (Fernandes and Carvalho, 2016; Thalhammer and Cingolani, 2014; Turrigiano, 2012). In vivo, global synaptic scaling takes place following sensory deprivation (Desai et al., 2002; Goel and Lee, 2007; Goel et al., 2006; Keck et al., 2013) or during sleep (Diering et al., 2017; De Vivo et al., 2017) and has been proposed to renormalize synaptic weights while maintaining the relative difference between incoming inputs and consolidating contextual memory. Uniform HSP could thus solve the apparent paradox of having circuits that are both stable and plastic (Davis, 2013; Galanis and Vlachos, 2020; Lee and Kirkwood, 2019; Turrigiano, 2017; Vitureira and Goda, 2013; Vitureira et al., 2012).

Yet, the multiplicative nature of synaptic scaling has recently been questioned (Hanes et al., 2020; Kim et al., 2012; Wang et al., 2019) and accumulating evidence shows that neurons implement a variety of homeostatic mechanisms acting on multiple spatial and temporal scales depending on activity perturbation paradigm, cell type, or developmental stage (Fernandes and Carvalho, 2016; Goel and Lee, 2007; Kim and Tsien, 2008; Lee and Kirkwood, 2019; Lee et al., 2014, 2013; Letellier et al., 2019; Queenan et al., 2012; Thiagarajan et al., 2005; Wierenga et al., 2006). For instance, excitatory synapses do not adapt uniformly in response to global activity deprivation involving the pharmacological blockade of AMPARs and/or NMDARs (Thiagarajan et al., 2005; Wang et al., 2019). Moreover, integrated and/or mature synaptic circuits appear less prone to undergo multiplicative scaling in comparison to immature circuits or *in vitro* models (Cingolani and Goda, 2008; Echegoyen et al., 2007; Goel and Lee, 2007; Hobbiss et al., 2018; Kim and Tsien, 2008; Lee and Kirkwood, 2019). Finally, altering the activity of individual connections revealed that subcellular compartments such as dendritic branches or individual synapses can sense local variations of activity and implement HSP in a relative autonomy (Barnes et al., 2017; Beique et al., 2011; Branco et al., 2008; Hou et al., 2008; Letellier et al., 2014, 2019; Li et al., 2018; Sutton et al., 2006).

One possible mechanism supporting such synapse autonomy is local protein translation, a process that occurs in remote subcellular compartments including presynaptic terminals and dendritic spines (Hafner et al., 2019; Rangaraju et al., 2017; Wang et al., 2010) and which contributes to local forms of HSP, in particular by regulating the expression of the GluA1 subunit of AMPARs (Aoto et al., 2008; Ju et al., 2004; Letellier et al., 2014; Maghsoodi et al., 2008; Sutton et al., 2006). Among the actors that can regulate local protein translation, microRNAs (miRNAs) control various forms of HSP (Dubes et al., 2019; Fiore et al., 2014; Letellier et al., 2014; Mellios et al., 2011; Rajman et al., 2017; Silva et al., 2019; Tognini et al., 2011). These small noncoding RNAs hybridize to the 3’ UTR of multiple target mRNAs and inhibit protein synthesis through translational repression or destabilization of the transcripts (Filipowicz et al., 2008; Friedman et al., 2008; Soula et al., 2018). In neurons, miRNAs can be found at the proximity of synapses where they respond to activity change and in turn regulate synaptic plasticity (Letellier et al., 2014; Park et al., 2019; Sambandan et al., 2017; Schratt, 2009).

miR-124 is one of the most enriched miRNAs in the brain and regulates synaptic function, including HSP, through the targeting of the GluA2 AMPAR subunit (Gascon et al., 2014; Ho et al., 2014; Hou et al., 2015). Specifically, the pharmacological blockade of action potentials (APs) and NMDARs in cultured neurons with TTX and D-APV, respectively, increases miR-124 expression and thus the repression of miR-124 on GluA2 translation which, unexpectedly, results in synaptic strengthening through the recruitment of GluA2-lacking AMPARs (Hou et al., 2015). Yet, there is also evidence that miR-124 elevation in vivo negatively regulates synaptic transmission and long-term potentiation (LTP) in the hippocampus with possible implications in spatial learning (Yang et al., 2012) and in neurological disorders such as epilepsy (Wang et al., 2016b), neurodegenerative diseases (Gascon et al., 2014; Wang et al., 2018) and multiple sclerosis (Dutta et al., 2013). Therefore, the mechanisms by which miR-124 regulates synaptic function still remain elusive. Another target of miR-124 is synaptopodin (SP) (Elramah et al., 2017), an actin-associated protein that controls HSP induced by activity deprivation with TTX or partial deafferentation of hippocampal granule cells (Vlachos et al., 2013). SP is an essential component of the spine apparatus, an endoplasmic reticulum-related organelle that is present only in a subset of dendritic spines where it regulates Ca^2+^ homeostasis and synaptic plasticity (Chirillo et al., 2019; Deller et al., 2000, 2003; Holbro et al., 2009; Korkotian and Segal, 2011; Vlachos et al., 2009, 2013). Intriguingly, SP clusters emerge inside spines with no obvious transport along dendrites, suggesting that the spine apparatus is assembled on site, possibly involving local protein translation (Konietzny et al., 2019). Yet, whether the local expression of SP is regulated by miR-124 and whether it provides autonomy to individual synapses to undergo HSP remains unknown.

Here, we tested the hypothesis that miR-124 controls HSP through the local coordination of SP and GluA2 translation. We report a synapse-autonomous mechanism for HSP, in which the ability of individual synapses to undergo HSP depends on the local expression of SP acting as a synaptic tag. Cultured hippocampal neurons respond to pharmacologically-induced activity deprivation by up-regulating AMPA receptors and SP in a non-uniform manner, resulting in a non-multiplicative increase of synaptic strengths. This process requires the de-repression of SP by miR-124 at large synapses which promotes the capture of surface diffusing AMPARs. By genetically silencing individual inputs in primary hippocampal neurons or organotypic slices, we further demonstrate that SP and AMPARs recruitment as well as spine growth operate in an input-specific manner. These findings uncover a miRNA-dependent and synapse-autonomous signaling in which a subset of synapses are primed for HSP.

## Results

### TTX-induced activity deprivation leads to the synaptic recruitment of AMPARs and SP in a non-uniform manner

We first checked whether chronic activity blockade of dissociated neurons using TTX altered the synaptic expression of AMPARs and SP (Gainey et al., 2009; Turrigiano et al., 1998; Vlachos et al., 2013; Wierenga et al., 2005). Because culturing rat hippocampal neurons in Neurobasal-containing medium occluded TTX-induced HSP in our conditions, most likely through inhibiting action potentials (APs) and spontaneous synaptic activity (**Fig. S1**), we opted for a culture medium that better fits physiological conditions and supports both neuronal activity and maturation, namely BrainPhys (Bardy et al., 2015; Satir et al., 2020) **(Fig. S1, see methods)**.

Hippocampal neurons cultured in this medium were transfected at DIV10 with Homer1c-GFP as a postsynaptic marker and immunostained at DIV15 for endogenous SP and surface AMPARs using an antibody raised against GluA2 AMPAR subunit that also recognizes GluA1 **(Fig. 1A, Fig. S1F and Fig. S2)**. Under basal conditions, ~25% of postsynapses contained SP clusters **(Fig. 1A,B)**, consistent with previous reports showing the accumulation of SP at the neck of a small fraction of spines in hippocampal pyramidal neurons (Deller et al., 2003; Orth et al., 2005; Vlachos et al., 2009). Synapses that were immunopositive for SP (SP+) were larger in size and displayed higher fluorescence intensity for immunostained AMPARs compared to synapses that were deprived of SP (SP-) **(Fig. 1C and Fig. S3A)**, indicating that the presence of SP is predictive of large and strong synapses (Vlachos et al., 2009). Upon 48 h TTX treatment, neurons displayed a higher percentage of SP+ synapses compared to untreated neurons, accompanied by an increase of synaptic AMPARs and Homer1c-GFP signals **(Fig. 1A-D and Fig. S3A,B)**. Interestingly, TTX treatment also enhanced the abundance of AMPARs selectively at the SP+ synapses, leaving SP-synapses with unchanged AMPAR content **(Fig. 1C)**. Consistent with this finding, TTX treatment induced a significant increase of immunostained AMPARs at large synapses (defined as Homer1c-GFP cluster area > 0.5 μm^2^) but not at small ones (Homer1c-GFP cluster area < 0.5 μm^2^) **(Fig. S3C,D)**. In addition, comparing the distribution of synaptic AMPAR fluorescence intensities revealed that the increase in AMPAR synaptic content observed for TTX-treated neurons compared to untreated ones was not multiplicative and selectively occurred at synapses with the highest AMPAR content **(Fig. 1E)**. Together, these results reveal a non-uniform synaptic recruitment of both SP and AMPARs across synapses during HSP and a selective contribution of the synapses displaying large size and high AMPAR content.

**Figure 1.**
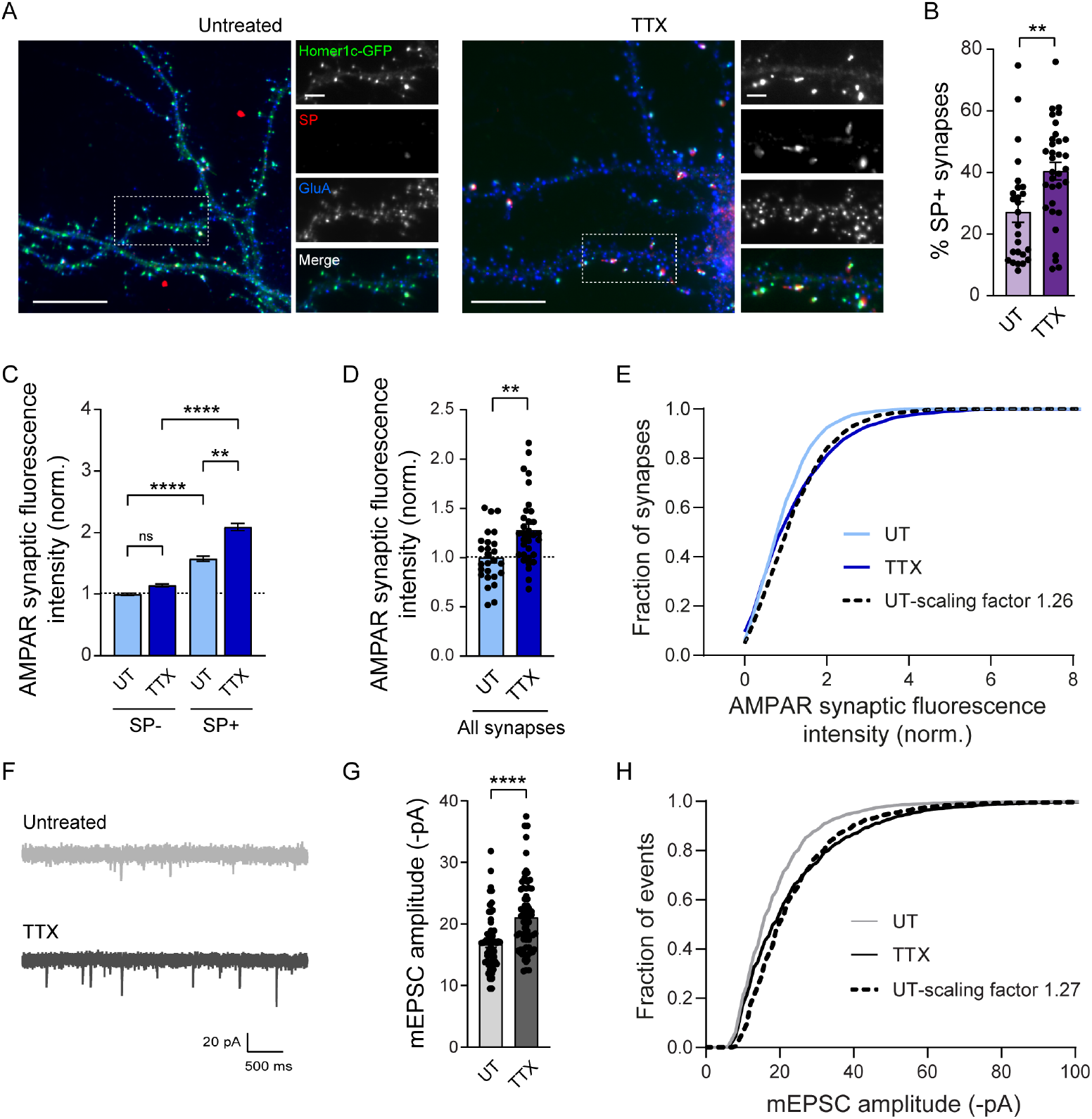
Cultured hippocampal neurons exhibit non-uniform homeostatic synaptic plasticity upon TTX treatment. **(A)** Micrographs showing Homerlc-GFP (green) and immunostaining for surface AMPARs (blue) and SP (red) in neurons treated with TTX, or untreated (UT). Scale bars, 20 μm (low magnification), 5 μm (insets). **(B)** Percentage of SP+ synapses for untreated (UT) and TTX-treated neurons (dot plots represent different cells). % SP+ spines: **P = 0.019 by Mann Whitney test. **(C)** AMPAR synaptic fluorescence intensity for SP+ versus SP-synapses in UT and TTX-treated neurons. AMPAR synaptic fluorescence intensity was normalized to SP- or small synapses, respectively, from UT condition. UT: SP-, n = 1455, SP +, n = 516; TTX: SP-, n = 1529, SP+, n = 724. n indicates the number of synapses. **P < 0.01, ****P < 0.0001, ns, not significant, P > 0.05 by Kruskal-Wallis test followed by Dunn’s multiple comparison test. **(D)** Average AMPAR synaptic fluorescence intensity in UT or TTX-treated neurons, regardless of the expression of SP (all synapses). AMPAR intensities were normalized to UT condition. **P = 0.0014 by Mann Whitney test. **(E)** Cumulative probability distribution of AMPAR synaptic fluorescence intensity for untreated neurons (UT, light blue), 48h TTX-treated neurons (TTX, dark blue) and scaled UT neurons (dotted black line, scaled factor = 1.26). UT vs TTX, P < 0.0001; UT-scaled vs TTX, P = 0.0217 by Wilcoxon matched-pairs signed rank test. **(F)** Representative traces of AMPAR-mediated miniature currents (mEPSCs) recorded from neurons cultured in BrainPhys™ supplemented with TTX for 48 h (TTX, dark gray), or left, untreated (UT, light gray). **(G)** Mean mEPSC amplitudes for each condition (UT: n = 65; TTX: n = 68). n indicates the number of cells. mEPSC amplitude: ****P < 0.0001 by Mann Whitney test. **(H)** Cumulative probability distributions of mEPSC amplitudes for untreated neurons (UT, light grey), TTX-treated neurons (TTX, black) and untreated scaled to TTX condition (dotted black line, scale factor = 1.27). The scale factor was obtained from the linear regression of the ranked mEPSC amplitudes for UT vs TTX condition. UT vs TTX, *P = 0.0412, UT scaled vs TTX, *P = 0.0316 by Kolmogorov-Smirnov test. Data represent mean ± SEM.

### TTX-induced activity deprivation leads to a non-multiplicative increase of synaptic strengths

To examine the functional correlate of these molecular changes observed by immunofluorescence, we performed patch-clamp recordings of AMPAR-mediated miniature currents (mEPSCs) in the same conditions. TTX-treated neurons exhibited larger mEPSCs (+4.4 pA) compared to untreated neurons **(Fig. 1F,G)**, confirming previous results (Sutton et al., 2006; Turrigiano et al., 1998; Vitureira et al., 2011). These currents displayed similar rise time and decay time constant as compared to the untreated condition **(Fig. S4A,B)** and were insensitive to a selective antagonist of Ca^2+^-permeable AMPARs, NASPM **(Fig. S4C-E)**. This suggested that AMPARs that were recruited at synapses upon TTX treatment contained the GluA2 subunit (Gainey et al., 2009). In addition, TTX treatment did not alter mEPSC frequency **(Fig. S1C,E and S4C,E)**, suggesting no change in the presynaptic release probability or number of active synapses. To investigate whether the increase in postsynaptic strengths was multiplicative, we scaled mEPSC amplitudes from control cells by the same factor (1.27) in order to match mEPSC amplitudes from TTX-treated cells. We next compared the cumulative distributions of scaled control mEPSC amplitudes and mEPSC amplitudes from TTX-treated cells and found a significant difference, indicating that HSP was not multiplicative **(Fig. 1H)**. Consistent with this finding, the rank-ordered mEPSC amplitudes from TTX-treated cells plotted against rank-ordered mEPSC amplitudes from control cells were better fitted with a quadratic function than with a linear function **(Fig. S4F)**. Together, these results suggest that large SP+ synapses are the ones displaying the largest increase in mEPSC amplitude upon TTX treatment.

### Surface-diffusing GluA2-containing AMPARs get immobilized at SP+ versus SP-synapses

To better understand how SP+ synapses get enriched in AMPARs relatively to SP-synapses, we next investigated the dynamics of surface GluA2-containing AMPARs at those two types of synapses by performing fluorescence recovery after photobleaching (FRAP) (Ashby et al., 2006). We expressed the GluA2 AMPAR subunit containing an N-terminal super-ecliptic (SEP) tag (SEP-GluA2) along with recombinant RFP-tagged SP (RFP-SP) in DIV8 cultured hippocampal neurons and performed live fluorescence imaging at DIV10. Under basal conditions, spines containing RFP-SP clusters exhibited a higher SEP-GluA2 intensity, compared to spines without RFP-SP **(Fig. 2A,B)**, demonstrating that recombinant SEP-GluA2 containing AMPARs behaved similarly to endogenous AMPARs, i.e. by accumulating preferentially at SP+ synapses. In dendritic spines lacking SP, SEP-GluA2 recovered from photobleaching with a time constant τ = 118.0 s; the recovery was still incomplete after 750 s with an immobile fraction of ~42% **(Fig. 2C,D)**, consistent with the synaptic turn-over of surface diffusing GluA2-containing AMPARs previously reported (Czöndör et al., 2012; Makino and Malinow, 2009; Penn et al., 2017). The recovery was lower for SP+ spines with a significantly larger immobile fraction of ~50% and a time constant τ = 128.3 s, suggesting that SP promotes the synaptic stabilization of surface diffusing AMPARs **(Fig. 2C,D)**. Together, these observations suggest that the recruitment of SP at a subset of synapses upon TTX treatment **(Fig. 1A,B)** contributes to the stabilization of surface-diffusing AMPARs during HSP **(Fig. 1C-E)**.

**Figure 2.**
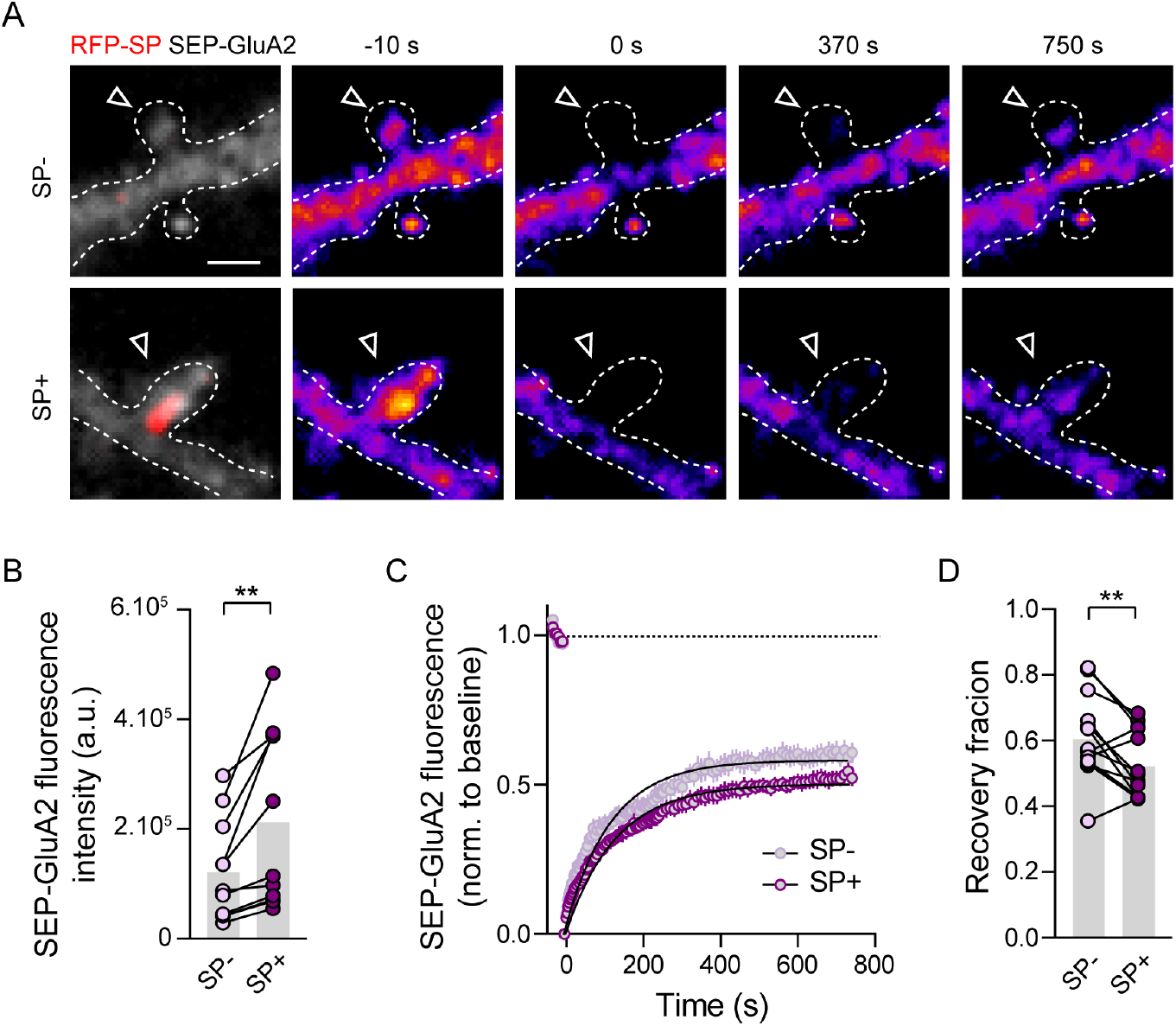
Higher immobilization of surface diffusing AMPARs at SP+ vs SP-synapses. **(A)** Example images of a FRAP experiment: SP- and SP+ spines from the same cultured hippocampal neuron transfected with SEP-GluA2 (gray and color-coded) + RFP-SP (red) 10s before, and 0, 370, 750 s after the photobleaching. **(B)** SEP-GluA2 fluorescence intensity at SP-vs SP+ spines. Each pair of dot plots represents average fluorescence for SP- and SP+ spines from a same neuron. **P = 0.0061 by two-tailed paired Student’s t test. **(C)** Quantification of FRAP dynamics for SP-vs SP+ spines. Recovery curves represent SEP-GluA2 fluorescence average per cell (n = 14 cells). The two traces were fitted using double exponential components equations and the convergence of the traces to a common fit was tested using the extra sum of squares F test. The F test indicates that the traces are best fitted by two divergent models (P < 0.0001). **(D)** Quantification of the recovery fraction 750 s after the photobleaching. Each pair of dot plots represents average recovery for SP- and SP+ spines from a same neuron. **P = 0.0067 by Wilcoxon matched-pairs signed rank test. Data represent mean ± SEM.

### SP is required for TTX-induced recruitment of synaptic AMPARs

To directly explore the role of endogenous SP in the recruitment of AMPARs during TTX-induced HSP, we next used a loss of function approach relying on the expression of a SP-targeting shRNA along with a GFP reporter (SP-shRNA-GFP). Using Western blotting, we found that SP-shRNA-GFP downregulates by ~30% recombinant RFP-SP expressed in COS-7 cells and this effect was rescued when expressing a shRNA-resistant RFP-SP construct, thus validating the specificity of the knock-down approach **(Fig. 3A,B)**. We then expressed SP-shRNA-GFP in DIV8 hippocampal neurons and performed immunostainings at DIV14-15. The immunofluorescence signal from endogenous SP was decreased by ~40% in neurons expressing SP-shRNA-GFP when compared to control neurons transfected with an empty vector **(Fig. 3C,D)**, validating both the specificity of the antibody against SP and the knock-down efficiency. Importantly, knocking-down endogenous SP in cultured neurons did not alter the basal synaptic accumulation of AMPARs, but inhibited the increase in synaptic AMPARs induced by TTX treatment **(Fig. 3E,F)**. Together with the observation that the presence of SP is predictive of large synapses and correlates with AMPAR stabilization **(Fig. 1C and Fig. S3A)**, these results indicate that the presence of SP is required for synapses to undergo HSP.

**Figure 3.**
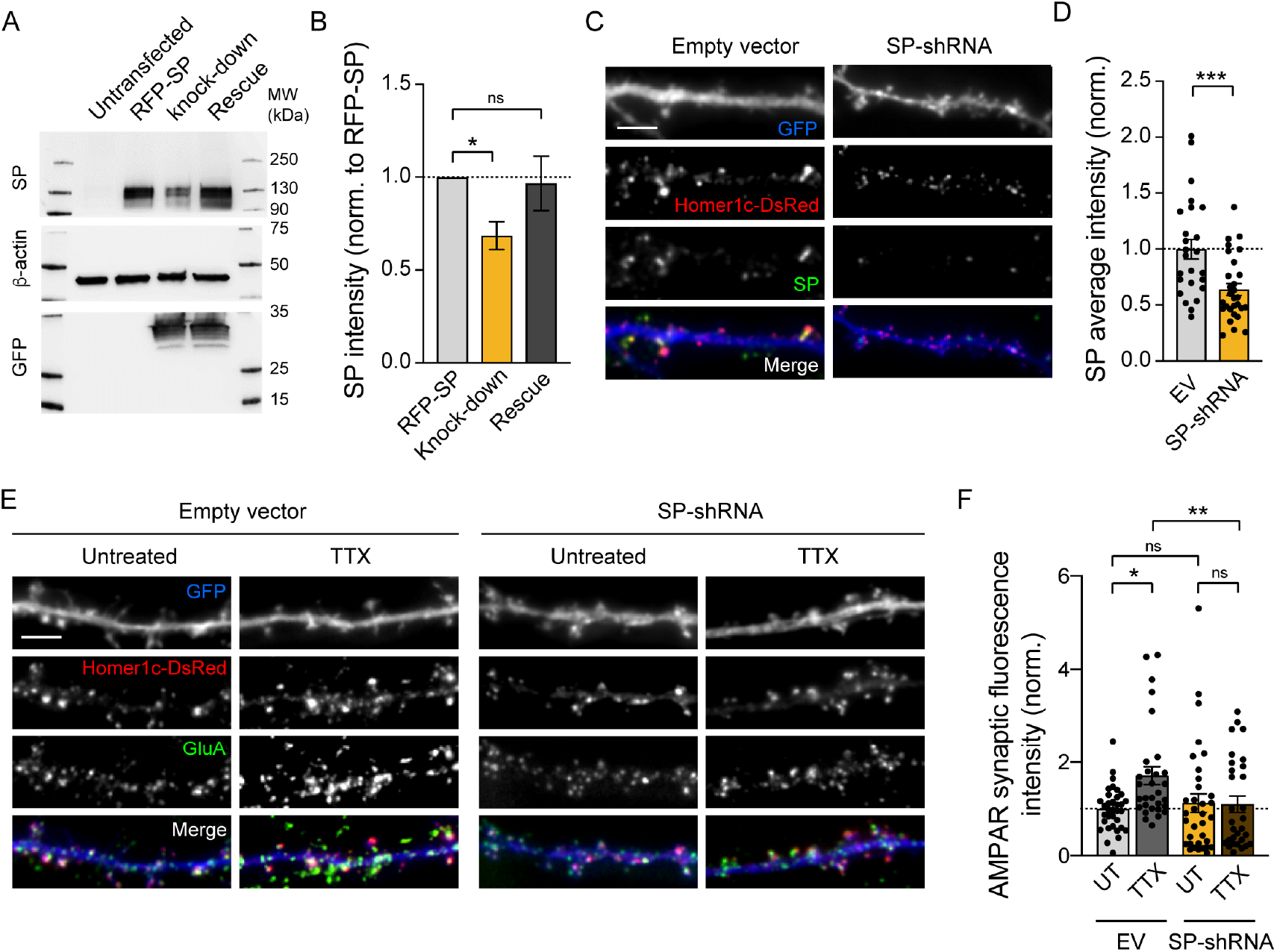
The synaptic recruitment of AMPARs induced by TTX treatment is controlled by SP. **(A)** Immunoblot analysis of SP in lysates from COS-7 cells which were untransfected, transfected with RFP-SP + empty vector or co-transfected with SP-shRNA-GFP + RFP-SP (knock-down) or shRNA-resistant RFP-SP (rescue). β-actin and GFP signals illustrate equal protein loading and SP-shRNA expression level, respectively. **(B)** Quantification of SP expression from 6 different experiments, normalized to the β-actin signal. *P < 0.05, ns, not significant, P > 0.05 by Kruskal-Wallis test followed by Dunn’s multiple comparison test. **(C)** Homer-DsRed (red) and immunostained endogenous SP (green) in dendrites from neurons transfected with either SP-shRNA or empty vector with GFP reporter (blue). Scale bar, 5 μm. **(D)** Average SP intensity for both conditions (EV: n = 25 cells; SP-shRNA: n = 29 cells; normalized to empty vector condition). ***P = 0.0007 by two-tailed Student’s t test. **(E)** Homer1c-DsRed (red) and immunostained surface AMPARs (green) in dendrites from neurons transfected with either SP-shRNA-GFP or empty vector (EV) with GFP reporter (blue) and treated with TTX or left untreated (UT). Scale bar, 5 μm. **(F)** AMPAR synaptic fluorescence intensity normalized to untreated empty vector (EV) condition (EV: UT, n = 42, TTX, n = 33; SP-shRNA: UT, n = 30, TTX, n = 31). n indicates the number of cells. *P < 0.05, **P < 0.01, ns, not significant, P > 0.05 by Kruskal-Wallis test followed by Dunn’s multiple comparison test. Data represent mean ± SEM.

### miR-124 is down-regulated in cultured hippocampal neurons upon prolonged TTX treatment

We next wondered whether the non-uniform upregulation of synaptic AMPAR and SP levels upon TTX treatment could result from the downregulation of miR-124, which is predicted to bind the 3’UTRs of both GluA2 and SP transcripts and thus might repress their expression in basal conditions (Elramah et al., 2017; Gascon et al., 2014; Ho et al., 2014). To test this hypothesis, we first investigated whether treating cultured hippocampal neurons with TTX could alter miR-124 expression levels as well as its targets *GluA2* and *synaptopodin* mRNAs by performing quantitative RT-PCR (qRT-PCR) in DIV15 neurons. The level of miR-124 was decreased by ~20% after 48 h TTX treatment compared to untreated neurons **(Fig. 4A)**. In contrast, the levels of two other miRNAs, miR-92a and miR-181, which target GluA1 and GluA2 3’UTRs, respectively, and whose expression are altered in response to different plasticity paradigms (Letellier et al., 2014; Saba et al., 2012; Sambandan et al., 2017) were not affected by TTX **(Fig. 4A)**. This revealed the selective contribution of miRNAs according to the activity-deprivation paradigm (Dubes et al., 2019). Importantly, the drop in miR-124 levels induced by the TTX treatment was compatible with the increased expression of SP and GluA2-containing AMPARs at a subset of spines. However, this was not accompanied by any change in the cellular levels of *GluA2* or *synaptopodin* mRNA **(Fig. 4B)**, suggesting that miR124 regulates SP or GluA2 expression in a local manner and/or by inhibiting the translation process.

**Figure 4.**
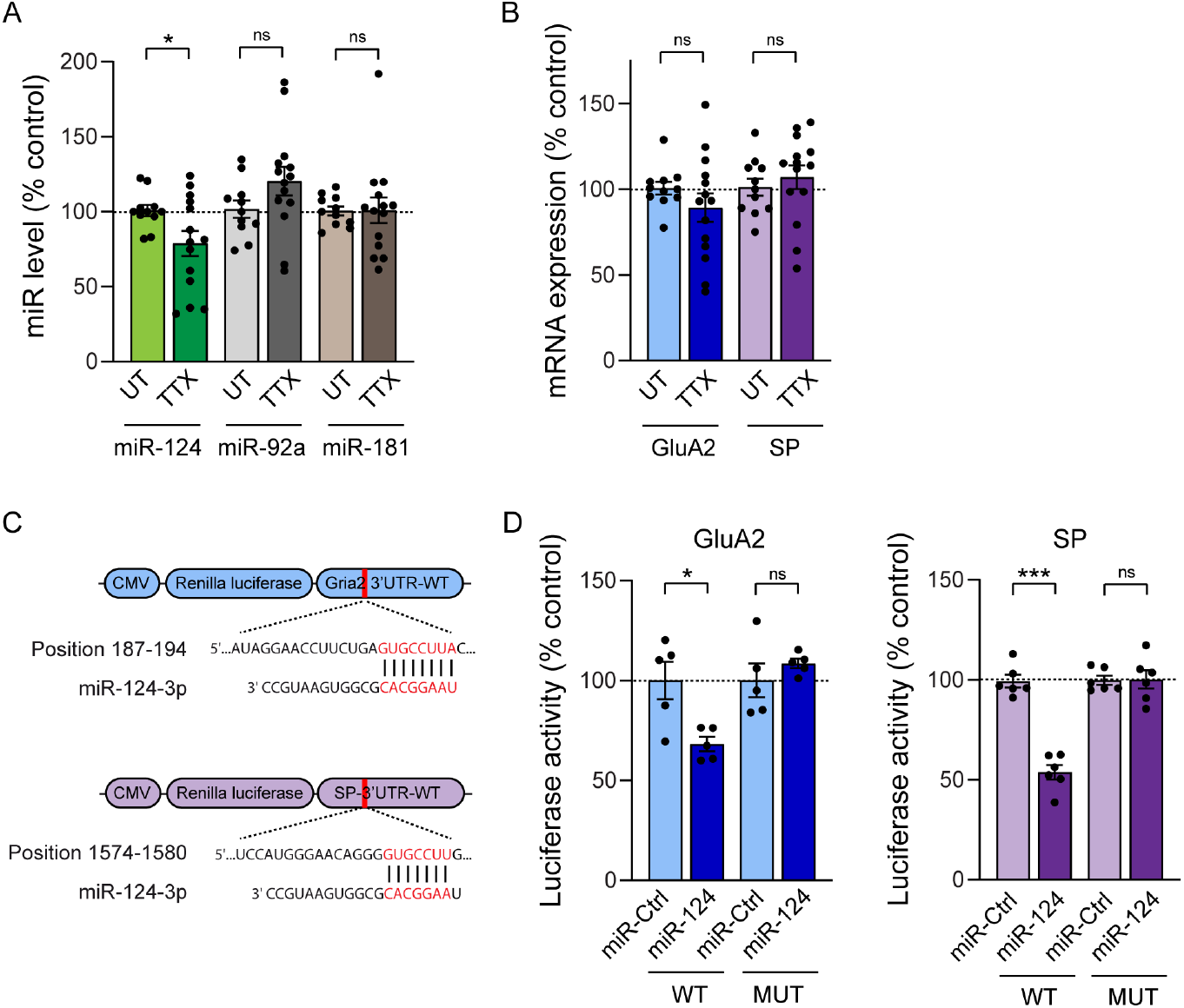
miR-124 level is selectively downregulated upon TTX treatment and directly inhibits GluA2 /SP translation. **(A,B)** Expression levels of miR-124, miR-92a and miR-181 (A), and GluA2 and SP mRNAs (B) determined by qRT-PCR in neurons treated with TTX or left, untreated (UT). Data are expressed as a percentage of the UT condition (miR-124: UT, n = 11, TTX, n = 14; miR-92a: UT, n = 11, TTX, n = 14; miR-181: UT, n = 11, TTX, n = 14; GluA2 mRNA: UT, n = 11, TTX, n = 14; SP mRNA: UT, n = 11, TTX, n = 14). *P < 0.05, ns, P > 0.05 by Mann Whitney test. n indicates the number of experiments. **(C)** Sequence alignment showing complementarity between Gria2 or SP 3’UTR and miR-124 binding seed region (highlighted in red). **(D)** Levels of luciferase activity measured from HEK-293 expressing miR-Ctrl or miR-124 together with the *Renilla luciferase* coding sequence reporter fused to the Gria2 or SP 3’UTR wild-type (WT) or mutated (MUT) to prevent miR-124 binding. Data are expressed as a percentage of miR-Ctrl condition (GluA2-3’UTR-WT: miR-Ctrl, n = 5, miR-124, n = 5; GluA2-3’UTR-MUT: miR-Ctrl, n = 5, miR-124, n = 5; SP-3’UTR-WT, miR-Ctrl, n = 6, miR-124, n = 6; SP-3’UTR-MUT: miR-Ctrl, n = 6, miR-124, n = 6). n indicates the number of experiments. *P < 0.05, ***P < 0.001, ns, not significant, P > 0.05 by Mann Whitney test. Data represent mean ± SEM.

### miR-124 inhibits translation of GluA2 and SP through direct interactions with their 3’UTR

To directly assess the ability of miR-124 to repress GluA2 and SP translation through binding to their respective 3’UTRs, we generated reporter plasmids by fusing the 3’UTR of *GluA2* or *SP* mRNA to the 3’ terminus of a *Renilla* luciferase coding sequence. We then co-expressed these constructs in HEK-293 cells together with miR-124 and measured luciferase activity **(Fig. 4C,D).** miR-124 expression significantly decreased the luciferase signal by ~30-45% for both constructs, compared to control miR-67 (miR-Ctrl) from *C. elegans* with no reported target in mammals **(Fig. 4D)**. Importantly, these effects were prevented when deleting miR-124 target regions in the 3’UTR of GluA2 and SP **(Fig. 4D)** indicating that miR-124 can inhibit both GluA2 and SP translation by directly interacting with their 3’UTR.

### miR-124 overexpression inhibits the synaptic recruitment of endogenous SP and the increase in synaptic strength upon TTX treatment

To examine whether miR-124 downregulation caused by TTX treatment was responsible for HSP, we next asked whether overexpressing miR-124 in cultured hippocampal neurons could impair the upregulation of endogenous synaptic AMPARs and SP induced by TTX treatment. In basal conditions, miR-124 overexpression did not significantly affect SP fluorescence intensity at synapses from DIV14 neurons, nor the percentage of SP+ synapses in comparison to miR-Ctrl **(Fig. 5A,B and Fig S5A)**. This suggested that endogenous miR-124 is highly expressed and already strongly represses SP expression in basal conditions. Moreover, synapse size, immunostained synaptic AMPARs or average AMPAR-mEPSC amplitude remained unchanged upon miR-124 overexpression **(Fig. 5C-F)**, which could be explained either by maximal repression of GluA2 synthesis by miR-124 under basal conditions or by the fact that miRNA-induced suppression of GluA2 expression is compensated by the increased assembly and synaptic recruitment of GluA2-lacking AMPARs at synapses (Hou et al., 2015; Silva et al., 2019). To test the latter hypothesis, we performed immunostaining of endogenous surface GluA1 using a specific antibody against the N-terminal part of GluA1 (Letellier et al., 2014) and found higher signal for neurons transfected with miR-124 in comparison to neurons transfected with miR-Ctrl **(Fig. S5C,D)**. Together with the lack of effect on AMPAR immunostaining and mEPSC amplitude, this observation suggests the molecular replacement of GluA2-containing AMPARs by GluA1-containing AMPARs in neurons overexpressing miR-124.

**Figure 5.**
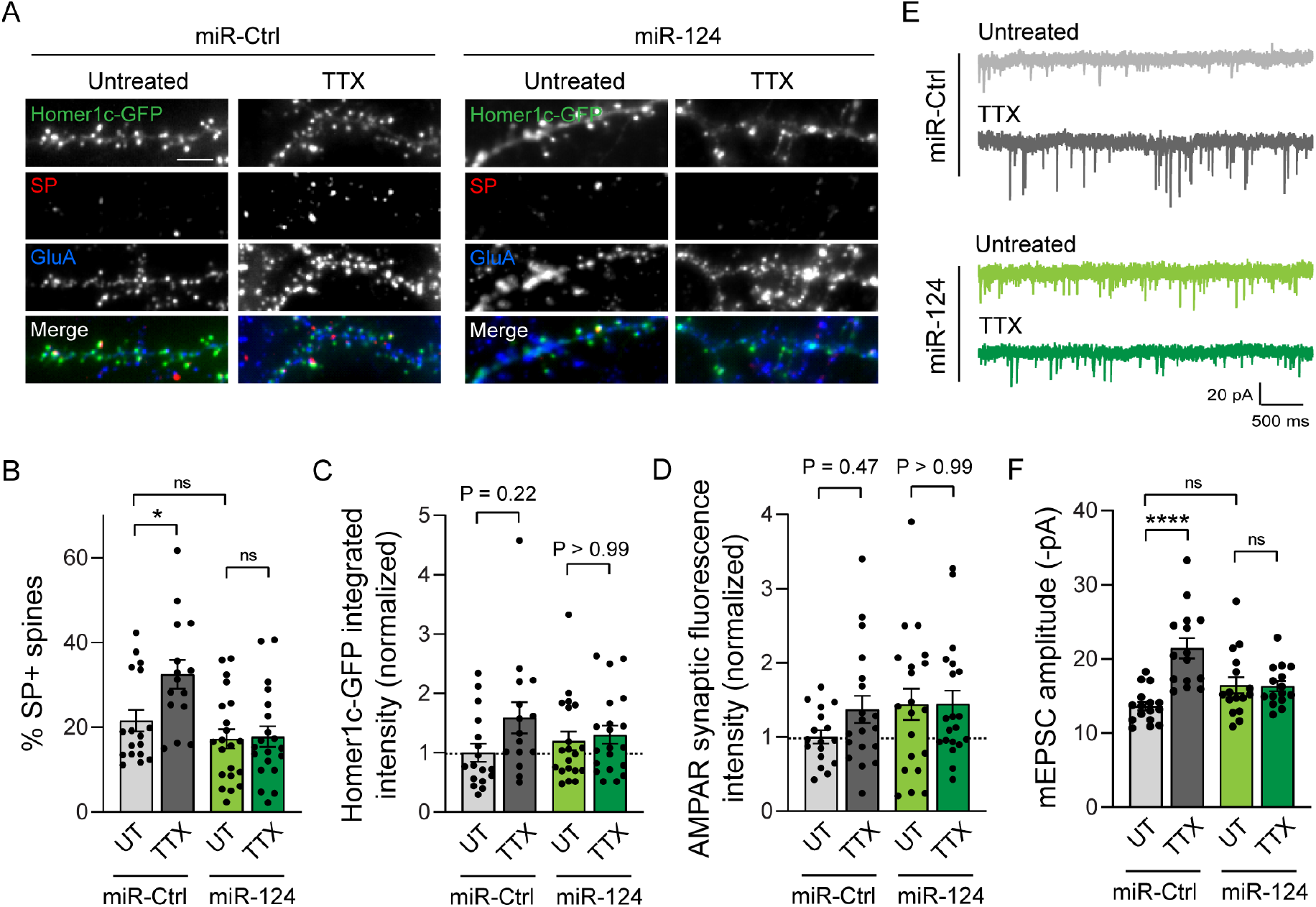
miR-124 overexpression inhibits TTX-induced HSP. **(A)** Micrographs showing Homer1c-GFP (green) and immunostaining for surface AMPARs (blue) and endogenous SP (red) in neurons transfected with miR-124 or control miR-67 (miR-Ctrl), and treated with TTX, or left untreated (UT). Scale bar, 5 μm. **(B)** Percentage of SP+ synapses for same conditions as in (A) (miR-Ctrl: UT, n = 17, TTX, n = 15; miR-124: UT, n = 20, TTX, n = 20). n indicates the number of cells. *P < 0.05, ns, not significant, P > 0.05 by two-way ANOVA test followed by Tukey’s multiple comparison test. **(C)** Homer1c-GFP intensity for same condition as in (A), normalized to untreated miR-Ctrl (miR-Ctrl: UT, n = 17, TTX, n = 15; miR-124: UT, n = 20, TTX, n = 20). n indicates the number of cells. ns, not significant, P > 0.05 by two-way ANOVA test. **(D)** AMPAR synaptic fluorescence intensity for same condition as in (A), normalized to untreated miR-Ctrl (miR-Ctrl: UT, n = 17, TTX, n = 15, miR-124: UT, n = 20, TTX, n = 20). n indicates the number of cells. ns, not significant, P > 0.05 by two-way ANOVA test. **(E)** Representative traces of AMPAR-mediated miniature currents (mEPSCs) recorded from neurons expressing miR-124 or miR-Ctrl in TTX-treated or untreated neurons. **(F)** AMPAR-mEPSC average amplitudes for same condition as in (E) (miR-Ctrl: UT, n = 16, TTX, n = 15, miR-124: UT, n = 16, TTX, n = 15). n indicates the number of cells. ****P < 0.0001, ns, not significant, P > 0.05 by Kruskal-Wallis test followed by Dunn’s multiple comparison test. Data represent mean ± SEM.

Importantly, overexpressing miR-124 blocked the significant increase in the percentage of SP+ synapses and average mEPSC amplitudes induced by TTX treatment as well as the trend for an increase in synapse size and synaptic AMPAR intensity **(Fig. 5A-F)**. These results suggest that the decrease of miR-124 level upon TTX treatment is required to enable the synaptic recruitment of AMPARs and SP as well as the increase in mEPSC amplitude upon TTX treatment. No significant change in mEPSC frequency was observed in any of the conditions **(Fig. S5B)**, indicating that miR-124 expression or TTX treatment have no effect on presynaptic function or the number of active synapses.

### The synaptic recruitment of SP upon TTX treatment requires the SP-3’UTR miR-124 binding region

We next investigated whether miR-124 binding to the SP 3’UTR could control the upregulation of SP in neurons during HSP. To test this hypothesis, we designed recombinant RFP-SP constructs fused to their respective 3’UTR sequences containing or lacking the miR-124 binding region (- WT or -MUT, respectively) and expressed them in cultured hippocampal neurons. This allowed us to selectively impair the interaction of endogenous miR-124 with recombinant SP transcripts without compromising the pathways mediated by other miR-124 targets. To limit the effect of overexpressing exogenous RFP-SP that could occlude the homeostatic response **(Fig. S6)**, we opted for a replacement strategy and co-expressed SP-shRNA with GFP reporter along with shRNA-resistant RFP-SP-3’UTR-WT/MUT constructs and Homer1c-BFP as a postsynaptic marker. In neurons expressing RFP-SP-3’UTR-WT, ~23% of synapses were found associated with RFP-SP clusters and this percentage reached ~33% after 48 h TTX treatment, thus reproducing the behavior of endogenous SP in control neurons and validating our replacement strategy **(Fig. 1A,B, Fig. 6A,B and Fig. S6)**. In contrast, neurons transfected with RFP-SP-3’UTR-MUT displayed ~35% of synapses containing RFP-SP clusters in basal conditions (**Fig. 6A,B)**. Importantly, this percentage did not further increase following TTX treatment, indicating that deleting the miR-124 binding region in the SP 3’UTR occluded the homeostatic response by de-repressing SP translation.

**Figure 6.**
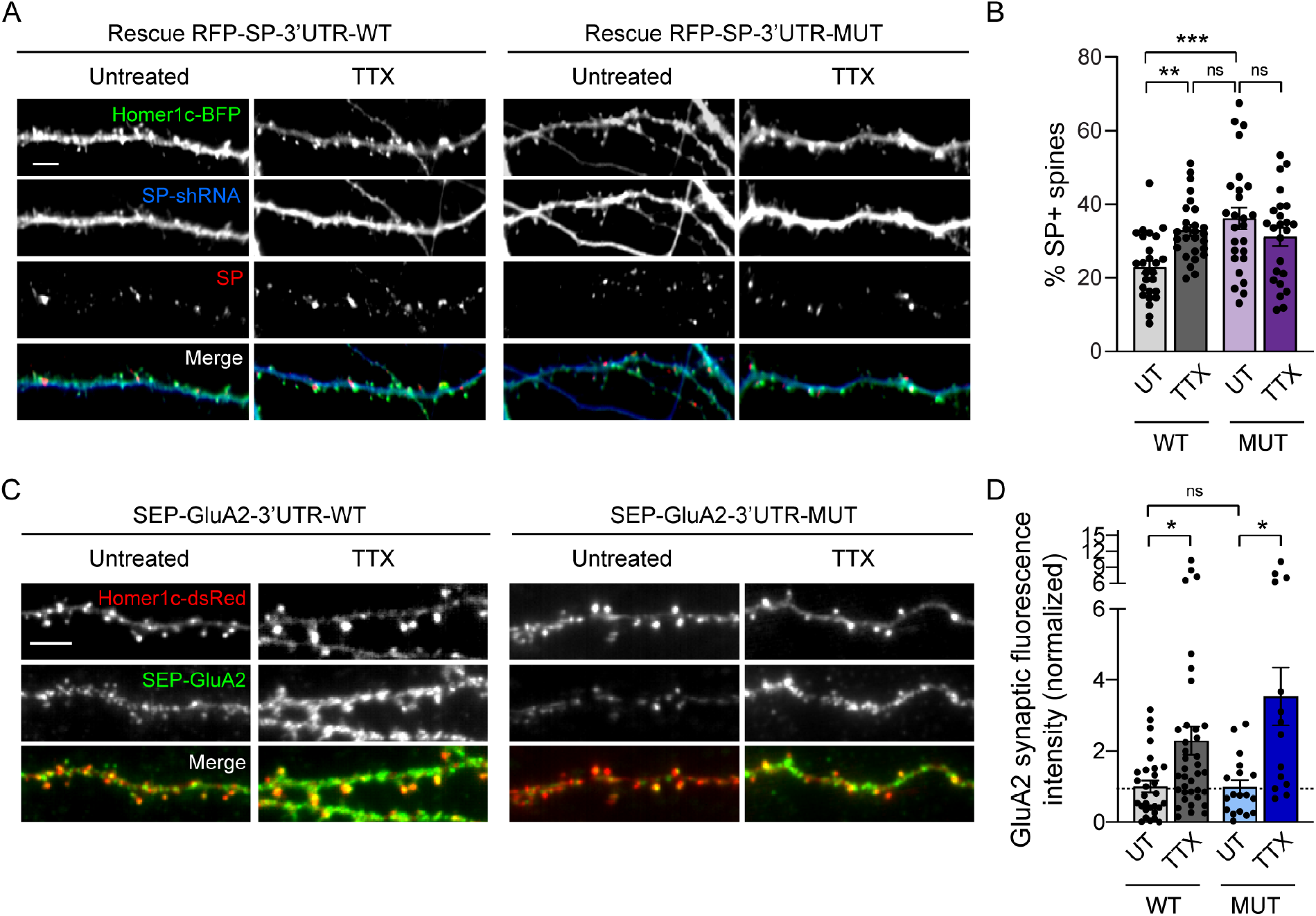
SP but not GluA2 synaptic recruitment upon TTX requires the 3’UTR miR-124 binding region. **(A)** Micrographs showing dendrites from neurons transfected with Homer1c-BFP (green), GFP-SP-shRNA (blue) and a rescue RFP-SP construct containing wild-type (WT) or mutated (MUT) 3’UTR (red) and treated with TTX, or left untreated (UT). Scale bar, 5 μm. **(B)** Percentage of SP+ synapses for each condition (3’UTR-WT: UT, n = 26, TTX, n = 28; 3’UTR-MUT: UT, n = 26, TTX, n = 24). n indicates the number of cells. **P < 0.01, ***P < 0.001, ns, not significant, P > 0.05 by two-way ANOVA test followed by Tukey’s multiple comparison test. **(C)** Micrographs showing dendrites from neurons transfected with Homer1c-DsRed (red) and SEP-GluA2 constructs containing wildtype (WT) or mutated (MUT) 3’UTR (green) and treated with TTX for 48 h, or left untreated (UT). Scale bar, 5 μm. **(D)** SEP-GluA2 synaptic fluorescence intensity for each condition, normalized to GluA2-3’UTR-WT untreated neurons (3’UTR-WT: UT, n = 29, TTX, n = 37; 3’UTR-MUT: UT, n = 18, TTX, n = 14). n indicates the number of cells. *P < 0.05, ns, not significant, P > 0.05 by Kruskal-Wallis test followed by Dunn’s multiple comparison test. Data represent mean ± SEM.

### The synaptic recruitment of GluA2-containing AMPARs upon TTX treatment does not require the GluA2-3’UTR miR-124 binding region

The de-repression of SP translation by miR-124 that occurs upon TTX treatment likely contributes to the synaptic stabilization of AMPARs at a new subset of synapses. We next wondered to what extent GluA2 expression could be similarly de-repressed upon TTX, and thus contribute to HSP by increasing the total pool of available AMPARs. Using a similar strategy as for SP, we examined the effect of expressing recombinant SEP-GluA2 constructs fused to 3’UTR sequences containing or lacking the miR-124 binding region (-WT or -MUT, respectively). Cultured neurons were transfected at DIV8 with SEP-GluA2-3’UTR-WT or SEP-GluA2-3’UTR-MUT and Homer1c-DsRed as a postsynaptic marker and subsequently processed at DIV14 for live immunostaining of surface recombinant AMPARs using an anti-GFP antibody **(Fig. 6C)**. In neurons transfected with SEP-GluA2-3’UTR-WT, TTX treatment induced a twofold increase in the synaptic accumulation of SEP-GluA2 compared to untreated neurons, showing that recombinant AMPARs contributed to HSP similarly to endogenous ones **(Fig. 6C,D)**. Surprisingly, however, deleting the miR-124 interacting region in the GluA2-3’UTR did not affect the basal surface levels of SEP-GluA2, nor its ability to get recruited at synapses upon TTX treatment **(Fig. 6C,D)**. These results suggest that endogenous miR-124 does not strongly repress GluA2 expression in basal conditions and make unlikely that GluA2 de-repression by miR-124 contributes to HSP.

### Sparse input silencing reveals synapse-autonomous recruitment of SP and AMPARs in cultured hippocampal neurons

Our results thus far suggest that miR-124 downregulation upon TTX treatment allows the de-repression of SP but not GluA2 expression at synapses. Importantly, the discrete distribution of SP at a subset of large synapses and the fact that TTX treatment increased the fraction of those synapses indicate that the homeostatic response is not uniform, possibly involving the local translation of SP (Cajigas et al., 2012; Konietzny et al., 2019; Yamazaki et al., 2008) under the control of miR-124. In support of this hypothesis, fluorescence *in situ hybridization* experiments using a specific anti-sense probe to visualize miR-124 revealed a clustered distribution of the fluorescence along dendrites of hippocampal neurons at DIV14 immunostained for pan-neuronal markers; the hybridization staining was absent at both somatic and dendritic levels when using a control scramble probe, demonstrating the specificity of the signal **(Fig. S7A,B)**.

To test for a local, synapse-autonomous, homeostatic upregulation of SP translation, we opted for a genetic approach where individual synaptic inputs are silenced through the expression of the tetanus toxin light chain (TetTX), which blocks neurotransmitter release through the proteolytic activity of the toxin against the requisite synaptic vesicle SNARE protein VAMP2 (Schiavo et al., 1992). We first examined in cultured hippocampal neurons whether the silencing of a subset of synaptic inputs using TetTx expression could induce a local homeostatic upregulation of AMPARs and SP at corresponding postsynapses. To this end, we carried out a sparse transfection of cultured neurons with a DNA construct in which the TetTx coding sequence was inserted downstream of the synaptophysin-GFP (Syp-GFP) sequence followed by the internal ribosome entry sequence (IRES) to visualize silenced presynaptic terminals with GFP (Ehlers et al., 2007; Harms et al., 2005). After 48 h of expression, the cultures were processed for immunostaining of surface endogenous AMPARs or endogenous SP and counterstained for MAP2 or PSD-95 to visualize dendrites or postsynaptic densities, respectively, and for VGLUT1 to visualize glutamatergic terminals. AMPARs and SP immunosignals in the postsynaptic neuron were higher when clusters were apposed to GFP positive boutons from transfected neurons compared to VGLUT1 immuno-positive boutons from non-transfected neurons **(Fig. S8A-D)**. This suggested that SP and AMPARs accumulate at silenced but not active synapses, thus representing a synapse-autonomous mechanism for HSP.

### Synapse-autonomous HSP requires SP-3’UTR miR-124 binding region at CA3 recurrent synapses from organotypic hippocampal slices

To test whether such synapse-autonomous HSP is also present in preserved neuronal circuits and depends on the interaction between miR-124 and SP-3’UTR, we turned to the CA3 recurrent circuit from organotypic hippocampal slices. We first sought to determine how SP distributes across spines from dendrites contacted by recurrent axon collaterals and whether its expression is regulated by miR-124. To this end, we transfected single CA3 pyramidal cells at DIV21 with RFP-SP-3’UTR-WT or -MUT through whole-cell patch-clamp recordings (Letellier et al., 2019) while knocking-down endogenous SP with SP-shRNA-BFP allowing to visualize dendritic spine morphology **(Fig. 7A)**. In neurons expressing RFP-SP-3’UTR-WT, we found that ~8% of dendritic spines contained SP-RFP clusters **(Fig. 7A,D)**. Those SP+ spines were larger compared to SP-spines, which was in agreement with our results in dissociated cultures and suggested that they contained more AMPARs **(Fig. 7A-C)** (Matsuzaki et al., 2001; Vlachos et al., 2009). Mutating SP-3’UTR to prevent miR-124 binding led to a threefold increase in the percentage of SP+ spines **(Fig. 7D)**. This effect was accompanied by a selective increase of size for the largest spines while the smallest spines remained of similar size **(Fig. 7E,F)**. Therefore, preventing SP regulation by miR-124 in organotypic slices mimicks the non-uniform regulation of synaptic strengths observed in primary neurons upon TTX treatment **(Fig. 1, S3 and S4)** and demonstrates that miR-124 exerts a continuous repression on SP translation in a subpopulation of spines.

**Figure 7.**
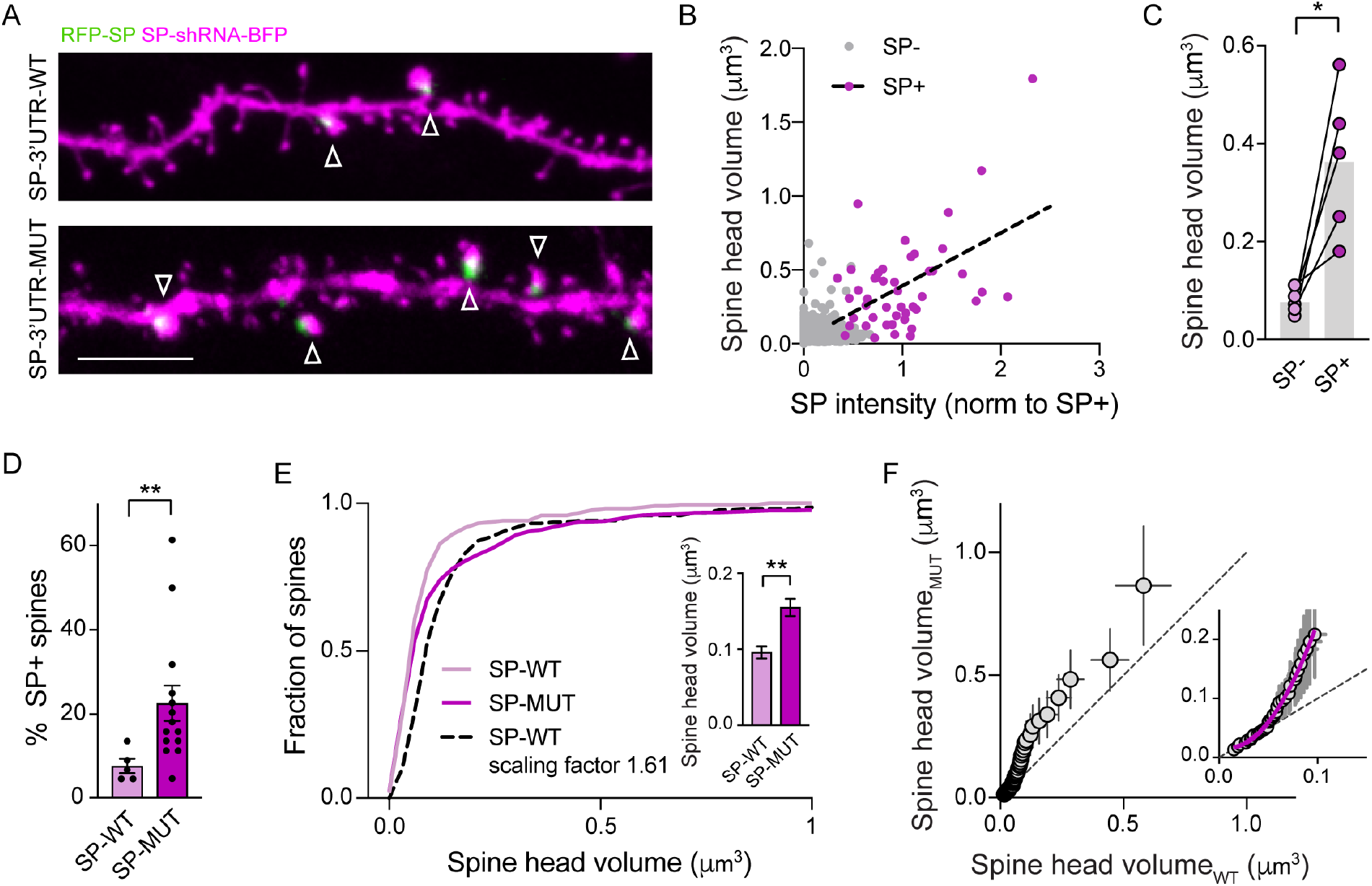
Non-uniform regulation of spine size by SP-3’UTR miR-124 binding region. **(A)** Confocal images showing dendrites from CA3 pyramidal cells transfected with RFP-SP (green) containing wildtype (WT) or mutated (MUT) 3’UTR + SP-shRNA-BFP (magenta). Arrowheads indicate SP+ spines. **(B)** Graph plot showing spine head volume vs SP integrated intensity (n = 5 cells). Each plot represents a single spine for which SP integrated intensity has been normalized to average SP intensity of SP+ spines in the same neuron. Plots for SP- and SP+ spines appear in grey and magenta, respectively. The black dotted line represents linear regression (R^2^ = 0.25, P = 0.0004). **(C)** Paired data showing spine head volume for SP-vs SP+ spines. Each pair of plot represents average values for SP-vs SP+ spines for a given neuron. *P = 0.0031 by two-tailed Student *t* test. **(D)** Percentage of SP+ spines in CA3 pyramidal cells transfected with SP SP-shRNA-BFP and rescue RFP-SP (green) containing wild-type (WT) vs mutated (MUT) 3’UTR (SP-3’UTR-WT: n = 5 cells; SP-3’UTR-MUT: n = 14 cells). **P = 0.0029 by Mann Whitney test. **(E)** Cumulative probability distributions of spine head volumes for neurons expressing RFP-SP-3’UTR-WT (SP-WT, light magenta) or RFP-SP-3’UTR-MUT (SP-MUT, dark magenta). The black dotted curve represents cumulative probability distribution corresponding to RFP-SP-3’UTR-WT condition scaled to RFP-SP-3’UTR-MUT (scale factor = 1.61). ****P < 0.0001 by Kolmogorov-Smirnov test. The inset shows average spine head volume for same conditions (SP-3’UTR-WT: n = 220; SP-3’UTR-MUT: n = 616). n indicates the number of spines. **P < 0.01 by Mann Whitney test. **(F)** Plot showing the rank-ordered spine head volumes for neurons expressing RFP-SP-3’UTR-WT vs RFP-SP-3’UTR-MUT. The rank-order plot was obtained by sorting from smallest to largest spine head volumes in SP-WT vs SP-MUT and plotting them against each other. The extra sum of squares F test indicates that the first values are better fitted with a second order polynomial quadratic curve (in magenta, SP-MUT = 0.02 – 0.42 x SP-WT + 25.32 x SP-WT^2^; R^2^ = 0.99; ****P < 0.0001) than with a linear regression (not shown, SP-MUT = – 0.05 + 2.42 x SP-WT).

We next sought to determine whether miR-124-dependent SP expression could be regulated upon local activity-deprivation. To do so, we took advantage of an approach where functionally connected CA3 pyramidal cells at DIV 21 are genetically manipulated through dual whole-cell patch-clamp recordings (Letellier et al., 2019). Using this strategy, presynaptic terminals from a single input were silenced through the whole-cell infusion of the GFP-syp-IRES-TetTx plasmid into the presynaptic neuron while endogenous SP was knocked-down in the postsynaptic cell through the infusion of SP-shRNA-EBFP and replaced by RFP-SP-3’UTR-WT or -MUT **(Fig. 8A-E)**. 48 h after transfecting neuron pairs through whole-cell patch-clamp, slices were fixed and processed for confocal microscopy. The occurrence of SP clusters as well as spine size were significantly higher for dendritic spines apposed to GFP-positive presynaptic terminals compared to neighboring spines from the same dendritic branch **(Fig. 8F-H)**. These results indicate that presynaptic silencing promotes the local recruitment of SP as well as spine growth. Importantly, this effect was partially occluded by the deletion of the miR-124 binding region in the SP 3’UTR, indicating the involvement of the de-repression of SP translation by miR-124 **(Fig. 8F-H)**.

**Figure 8.**
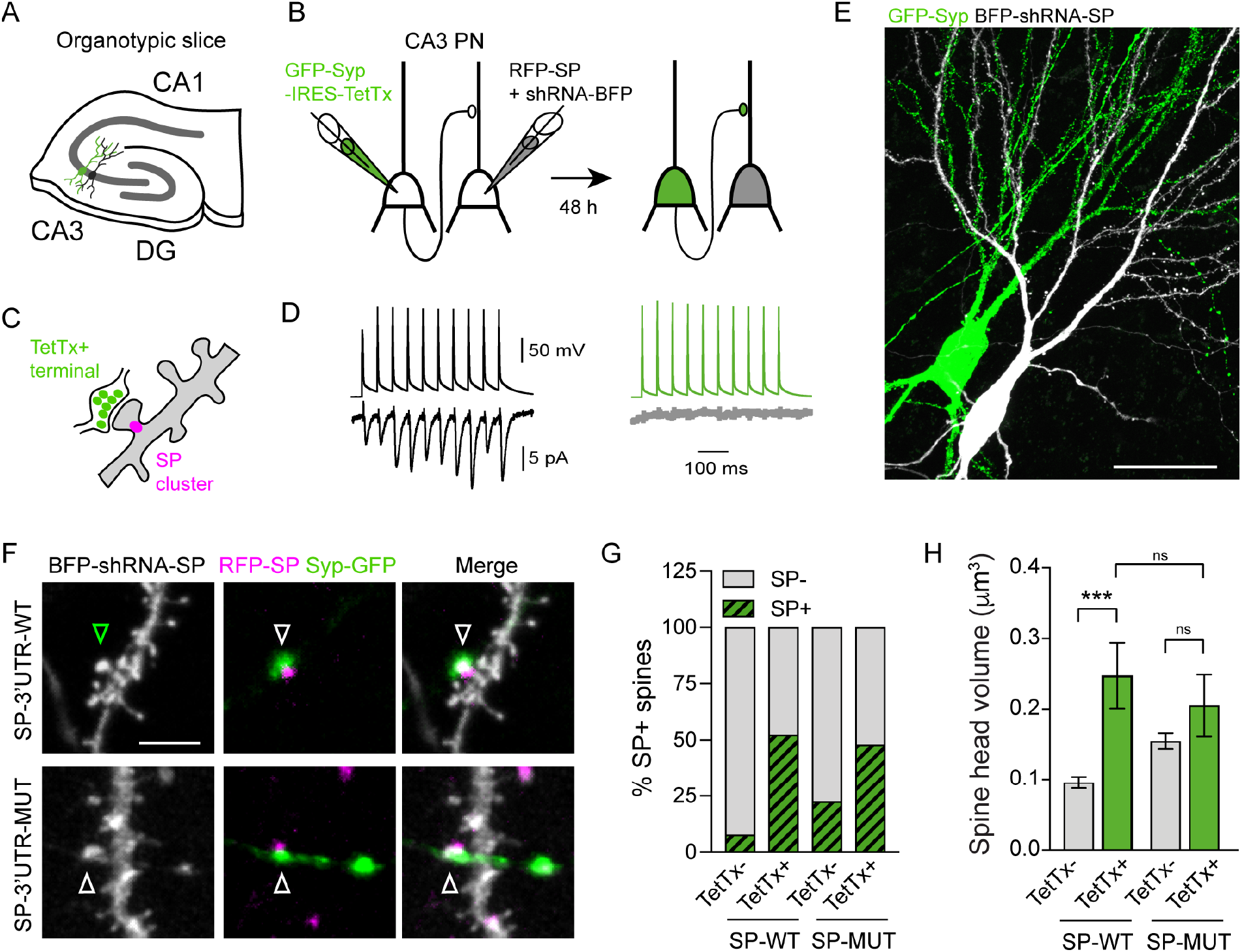
miR-124-dependent synapse-autonomous HSP at CA3 recurrent synapses. **(A)** Experimental design to silence individual presynaptic inputs in recurrent circuits between CA3 pyramidal cells in organotypic hippocampal slices. **(B,C)** Whole-cell recordings are used to ensure the functional connectivity between two CA3 pyramidal cells while infusing plasmids encoding GFP-Syp-IRES-TetTx and RFP-SP + BFP-shRNA-SP in the pre- and postsynaptic cells, respectively (B). Slices were fixed 48 h after transfection and processed for confocal microscopy in order to analyze spines contacted by GFP+ terminals vs neighboring spines from the same dendritic branch (C). **(D)** Pair recordings from functionally connected neurons during transfection (left) and 24 h after transfection (right). Top traces (black and green) show train of 10 action potentials elicited in the presynaptic cell. Bottom traces show corresponding evoked postsynaptic currents (black, average from 20 sweeps) during transfection but not 24 h after (grey, average from 20 sweeps). **(E)** Confocal image showing a pair of pre and postsynaptic CA3 pyramidal neurons transfected with GFP-Syp-IRES-TetTx (green) and RFP-SP (not shown) + SP-shRNA-BFP (grey), respectively. **(F)** Higher magnification of dendritic spines from neurons transfected with RFP-SP (magenta) containing wild-type (WT) or mutated (MUT) 3’UTR and BFP-shRNA-SP (grey) and contacted by GFP-Syp+ terminals (green). Arrowheads indicate putative synaptic contacts. **(G)** Percentage of SP+ spines in neurons expressing RFP-SP-3’UTR-WT vs -MUT and depending on the apposition or not to a GFP-Syp+ terminal (SP-3’UTR-WT: TetTx-, n = 220, TetTx+, n = 26; SP-3’UTR-MUT: TetTx-, n = 616, TetTx+, n = 25). **(H)** Spine head volume in same conditions as in (G) (SP-3’UTR-WT: TetTx-, n = 220, TetTx+, n = 26; SP-3’UTR-MUT: TetTx-, n = 616, TetTx+, n = 25). n indicates the number of spines. ***P < 0.001, ns, not significant, by Kruskall-Wallis test followed by Dunn’s multiple test. Data represent mean ± SEM.

## Discussion

We found an unexpected synaptic tagging mechanism for HSP by which the ability of individual synapses to respond to global or input-specific activity-deprivation depends on the local expression of SP, which promotes AMPARs recruitment and spine growth **(Fig. S9)**. Specifically, treating hippocampal neurons with TTX decreases miR-124 cellular levels, correlated with a non-uniform recruitment of SP and AMPARs across synapses and a non-multiplicative increase of synaptic strengths. By expressing miR-124, shRNAs to SP, or recombinant SP and GluA2 bearing mutated 3’UTR to prevent their targeting by native miR-124, we show that the homeostatic response requires the derepression of SP, but not AMPARs, induced by TTX. Finally, by genetically silencing individual inputs with tetanus toxin in primary cells or CA3 recurrent circuits from organotypic slices, we show that HSP involving SP up-regulation is synapse-autonomous and requires the 3’UTR miR-124 binding region.

### Multiplicative versus non-uniform HSP

Since the princeps studies in cultured cortical neurons (O’Brien et al., 1998; Turrigiano, 2008; Turrigiano et al., 1998), the prevailing view has been that postsynaptic strengths are uniformly scaled by a same factor in response to global change of network activity, a phenomenon that has been proposed to take place *in vivo* early during development following sensory deprivation (Desai et al., 2002; Goel and Lee, 2007; Goel et al., 2006; Keck et al., 2013). However, we report here in cultured hippocampal neurons that, in response to 48 h TTX incubation, relatively large and strong synapses scale with a higher gain compared to small and weak synapses. Our results recapitulate previous findings obtained in vitro or in vivo showing a “divergent scaling” of synaptic strengths (Echegoyen et al., 2007; Hanes et al., 2020; Lee et al., 2013; Thiagarajan et al., 2005) or spine sizes (Hobbiss et al., 2018) upon activity deprivation, where synaptic gain increases with initial postsynaptic strength or size. Besides the possible bias introduced by the method analysis used to test the multiplicative nature of HSP (Hanes et al., 2020; Kim et al., 2012), the discrepancy across studies might result from differences in experimental conditions, including the developmental stage (Goel and Lee, 2007), the duration of the activity perturbation (Hobbiss et al., 2018) and the culture conditions (Bardy et al., 2015; Ivenshitz and Segal, 2010).

The non-multiplicative HSP that we describe here, by selectively strengthening the strongest synapses may serve to restore normal firing rate in an optimized way (Hanes et al., 2020). However, by enhancing relative differences between postsynaptic strengths rather than scaling them uniformly, this form of HSP likely interferes with prior Hebbian synaptic changes and is expected to affect the relative capacity of large versus small synapses to undergo subsequent LTP (Thiagarajan et al., 2007). Accordingly, it was recently shown in organotypic slices that TTX-induced HSP prevents scaled synapses to undergo future LTP by altering short-term glutamate release dynamics (Soares et al., 2017) while reducing the threshold for postsynaptic LTP at small synapses (Hobbiss et al., 2018). Together, these results suggest that HSP induced by activity-deprivation at the network level alters synaptic strengths distribution and profoundly affects the rules for individual synapses to undergo Hebbian plasticity and to functionally interact together (Lee and Kirkwood, 2019), with possible consequences for synaptic circuit development (Tien and Kerschensteiner, 2018) and the formation of memory engrams (Lee and Kirkwood, 2019; Mendez et al., 2018). In particular, non-uniform HSP resulting from network-wide activity alterations could become maladaptive in some pathological contexts such as Alzheimer disease and epilepsy where it may be achieved at the expense of synaptic input integration and plasticity (Galanis and Vlachos, 2020; Li et al., 2019; Lignani et al., 2020; Styr and Slutsky, 2018).

#### SP as a molecular tag to promote input-specific synaptic plasticity

We provide a molecular mechanism for the non-uniform and synapse-autonomous HSP that is induced by activity deprivation. Specifically, we demonstrate that the divergent behavior of synapses facing local or global activity-deprivation can be attributed to the non-uniform distribution of SP, an essential component of the smooth ER-derived organelle called spine apparatus (Deller et al., 2003) which associates with a subpopulation of large and strong synapses (Orth et al., 2005; Spacek and Harris, 1997). Our immunocytochemistry data suggest that the increased expression of SP at a subset of synapses following activity-deprivation with TTX is responsible for the recruitment and stabilization of AMPARs. Our FRAP experiments further indicate that AMPARs stabilization at SP+ synapses occurs through the alteration of lateral diffusion, as previously evidenced for mGluR5 (Wang et al., 2016a); this may involve calcium release from internal stores (Vlachos et al., 2009) and/or actin remodeling events (Konietzny et al., 2019; Wang et al., 2016a). Whatsoever, the role of SP in AMPARs synaptic stabilization is activity-dependent since SP knock-down or knock-out strongly impairs both HSP and LTP but not average basal synaptic properties (Deller et al., 2003; Vlachos et al., 2013). In addition, our experiments in primary cultures and organotypic slices indicate that SP cluster formation, AMPARs synaptic accumulation and spine growth can be stimulated at individual synapses in response to the blockade of presynaptic glutamate release through TetTx expression. In agreement with our results, a recent electron microscopy study performed in the intact hippocampus revealed that mushroom spines but not smaller protrusions (e.g. thin, stubby or filopodial) exhibit higher volume and PSD size when contacted by TetTx+ presynaptic terminals and more likely contained a spine apparatus (Zhu et al., 2021). The input-specificity of SP recruitment upon activity perturbation is also supported by previous reports showing that high-frequency stimulation in the dentate gyrus enhances SP immunoreactivity selectively in the corresponding stimulated layers (de Solis et al., 2017; Yamazaki et al., 2008). Such synapse-specific regulations are in agreement with the non-uniform distribution of SP, which likely serves as a molecular tag enabling individual synapses to undergo potentiation. The appearance of SP clusters likely reflects the elaboration and the stabilization of the spine apparatus (Perez-Alvarez et al., 2020) that is, together with the presence of polyribosomes, predictive of activity-dependent spine enlargement (Chirillo et al., 2019). The fact that SP enables individual synapses to undergo potentiation in both Hebbian and homeostatic forms of plasticity underscores its fundamental role and raises the possibility that both types of plasticity coexist and/or occlude each other at individual synapses, possibly reflecting a shift in homeostatic setpoint and/or LTP sliding threshold (Galanis and Vlachos, 2020; Lee and Kirkwood, 2019; Li et al., 2019).

The fact that individual synapses can homeostatically adjust their own strength has also been reported previously using sparse expression of the potassium channel kir2.1 across presynaptic terminals to decrease the firing of individual inputs. Similar to our findings, the input-specific decrease of activity using Kir2.1 leads to the accumulation of AMPARs at corresponding spines but not neighboring ones, through a mechanism involving the downregulation of the activity regulated cytoskeleton-associated protein Arc (Beique et al., 2011; Hou et al., 2008). Arc is a protein involved in long-term synaptic depression and inverse synaptic tagging and can be locally synthesized (Bagni et al., 2000; El-Boustani et al., 2018; Nikolaienko et al., 2018; Okuno et al., 2012; Yin et al., 2002). However, in this paradigm, HSP involves the recruitment of GluA2-lacking AMPARs and is not supported by an increase in spine head volume, suggesting that the underlying mechanisms are different (Beique et al., 2011). Whether this form of synapse-specific HSP is regulated by miRNAs has not been investigated yet.

#### Synapse-autonomous regulation of SP but not GluA2 by miR-124

What are the mechanisms regulating SP recruitment in response to synapse-specific activity alteration? We demonstrate that the appearance of SP clusters at synapses following chronic activity deprivation depends on the presence of the miR-124 binding region in the 3’UTR of SP mRNA. Specifically, deleting the miR-124 interaction region in SP-3’UTR occludes the increased expression of SP at a subset of synapses that is induced by activity deprivation, revealing that miR-124 exerts a continuous repression on SP expression in some synapses and that this repression is released when activity is suppressed. Together with the fact that (1) this mechanism can be implemented in a synapse-specific manner, (2) SP mRNA and miR-124 are localized in the dendritic compartment according to *in situ* hybridization or trancriptomics experiments (this study, Cajigas et al., 2012; Ho et al., 2014; de Solis et al., 2017) and (3) SP clusters are directly assembled in spines (Konietzny et al., 2019), these observations indicate that SP synthesis can be locally stimulated through the de-repression by miR-124 to support input-specific increase of synaptic strength. The role of miR-124 downregulation in activity-dependent synaptic potentiation is further evidenced by the fact that miR-124 overexpression inhibits synaptic strength increase upon TTX treatment (our study) or after eliciting LTP (Yang et al., 2012) while miR-124 neutralization using a sponge promotes AMPARs synaptic recruitment on its own (Hou et al., 2015).

In contrast to SP, deleting the miR-124 interaction region in the GluA2 3’UTR alone is not sufficient to increase the basal expression of AMPARs at synapses, or their recruitment upon TTX treatment. This might be explained by the fact that additional mechanisms are required to stabilize AMPARs at synapses, among which the derepression of SP synthesis by miR-124, as discussed above. It is also possible that GluA2 basal expression is not strongly repressed by miR-124 at the proximity of synapses; this is suggested by ISH experiments showing low abundance of GluA2 transcripts in the synapto-dendritic compartment (Ho et al., 2014) and by the fact that GluA2 synaptic expression can be repressed when overexpressing miR-124 (Gascon et al., 2014; Ho et al., 2014; Hou et al., 2015). In one possible model, GluA2 is primarily synthesized at the cell body level and subsequently targeted to activity-deprived synapses through trafficking mechanisms (Chater and Goda, 2014; Hangen et al., 2018), in which SP and the spine apparatus participate.

While the implication of miR-124 downregulation in our HSP paradigm does not seem to rely on GluA2 synthesis, the situation might be different in other activity-deprivation paradigms involving different miRNAs. In particular, miR-186-5p exerts a continuous repression on GluA2 synthesis in primary hippocampal neurons, that is released upon AMPARs and NMDARs pharmacological blockade (Silva et al., 2019). Interestingly, this paradigm affects neither miR-124 nor miR-92a global levels, and rather results in a uniform upscaling of synaptic strengths. Therefore, the control of GluA2 expression by miR-186-5p occurs at the cell body level and affect synapses more widely and uniformly than following APs blockade with TTX. Altogether, these findings underscore the existence of multiple miRNAs dependent pathways that neurons can use to adjust HSP in time and space depending on the type of activity perturbations (Dubes et al., 2019).

#### miR-124, a versatile miRNA controlling various types of synaptic plasticity

Several lines of evidence indicate that miR-124 can regulate synaptic function through other targets than SP. In particular, it was previously reported that miR-124 elevation can also drive HSP by directly repressing GluA2 expression, when activity deprivation is induced by inhibiting both APs and NMDARs with TTX and AP5, respectively (Hou et al., 2015). In this case, the repression of GluA2 synthesis by miR-124 promotes the synaptic insertion of homomeric GluA1-containing AMPARs, and may act in conjunction with the de-repression of GluA1 synthesis by miR-92a (Letellier et al., 2014). The fact that miR-124 elevation favors the assembly and insertion of GluA2-lacking AMPARs is also supported by our experiments where overexpressing miR-124 induces a synaptic increase of the GluA1 AMPAR subunit, although we found that this effect is not accompanied by an increase in synaptic strength. Therefore, the bidirectional regulation of miR-124 expression, which likely depends on NMDARs activity and the duration of activity deprivation, may represent a functional switch to produce a selective homeostatic response with respect to the AMPARs subunit composition that confers specific plastic properties to synapses (Diering and Huganir, 2018; Gascon et al., 2014). In addition to the role of miR-124 in HSP through the targeting of SP or GluA2, there is also evidence that miR-124 negatively regulates synaptic transmission, LTP and spine density through the targeting of tyrosine-protein phosphatase non-receptor type 1 (PTPN1) that regulates GluA2 synaptic insertion (Wang et al., 2018), and through the targeting of the transcription factors Zif268 (Yang et al., 2012) and CREB1 (Wang et al., 2016b), with possible implications in spatial learning, epilepsy and Alzheimer disease. In one possibility, the de-repression of these pathways caused by miR-124 downregulation could contribute to the homeostatic increase of synaptic strengths that we describe here. Considering the implication of miR-124 and SP in both homeostatic and Hebbian plasticity and the fact that these two types of plasticity exhibit synapse-autonomous features, it will be important to investigate how the multiple miR-124-dependent pathways cooperate in time and space to control the interplay between Hebbian and homeostatic plasticity at individual synapses and how this balance is disrupted in pathological conditions.

## Acknowledgments

We acknowledge S. Okabe, D. Choquet, D. Perrais and A. Triller/C. Specht for the generous gift of plasmids, D. Choquet and C. Breillat for the anti-GluA1 antibody, F. Levet and J-B. Sibarita for support using SpineJ plugin. We thank M. Munier for expert technical assistance in molecular biology and preparation of cultures. We also thank the Bordeaux Imaging Center, part of the France BioImaging national infrastructure ANR-10-INBS-04S for support in confocal microscopy, the animal facility of the University of Bordeaux (in particular M. Deshors, S. Pavelot, C. Martin and P. Costet), the cell culture facility of the Interdisciplinary Institute for Neuroscience (in particular N. Retailleau, E. Verdier), R. Sterling, J. Carrere, and J. Girard for technical assistance. The protein quantification was done in the Biochemistry and Biophysics Core Facility of the Bordeaux Neurocampus (Bioprot) with the help of Yann Rufin. This work received funding from the Centre National de la Recherche Scientifique, Agence Nationale pour la Recherche (grant “SynSpe” ANR-13-PDOC-0012-01), Investissement d’Avenir Labex BRAIN, and Fondation pour la Recherche Médicale (“Equipe FRM” DEQ20160334916).

## Author contributions

Conceptualization, M.L., O.T., A.F. and S.D.; Methodology, M.L., O.T., A.F., B.T., S.B. and C.P.; Investigation, S.D., M.L., A.S., S.B. and B.T.; Data curation, S.D., M.L., B.T. and A.S.; Formal Analysis, S.D. and M.L.; Writing - original draft, M.L.; Writing - Review and Editing, M.L., O.T., S.D. and A.F.; Visualization, S.D. and M.L.; Funding Acquisition, M.L. and O.T.; Supervision, M.L. and O.T.; Project Administration, M.L. and O.T.

## Experimental procedures

### DNA plasmids

The pcDNA Homer1c-GFP was gift from S. Okabe (Tokyo University, Japan). The pcDNA Homer1c-DsRed and SEP-GluA2 3’UTR plasmids was already described (Mondin et al., 2011). To generate SEP-GluA2-3’UTR-MUT, we inserted the mutated sequence (synthetized by Eurofins) at the HpaI/AflII sites in place of the WT sequence. HA-GluA1 and HA-GluA2 plasmids were gift from D. Choquet (Interdisciplinary Institute for Neuroscience, Bordeaux). RFP Synaptopodin was a gift from A. Triller (École Normale Supérieure, Institut de Biologie de l’ENS, Paris). RFP-SP-3’UTR-WT / -MUT were generated by inserting the partial WT or mutated 3’UTR sequence of SP (GTCTCCATGGGAACAGGGGTGCCTTGTCAGTG or GTCTCCATGGGAACAGGGCACGGAAGTCAGTG respectively) at ApaI / MluI sites in the RFP-SP plasmid. The corresponding rescue forms Rescue RFP-SP-3’UTR-WT / -MUT were generated by inserting the sequence resistant to the sh (synthetized by Eurofins) at the XbaI / BamHI sites. EGFP-synaptophysin:IRES:TetTx construct was a gift from D. Choquet / D. Perrais (Interdisciplinary Institute for Neuroscience, Bordeaux). The target sequence of the shRNA against SP (SP-shRNA with GFP reporter) was designed with BLOCK-it RNAi designer (https://rnaidesigner.thermofisher.com/rnaiexpress/), the selected sequence (5’-GGTGTATAGTGAAGTACATCT-3’) was inserted in a miR-30 context (pCAG-miR-30, from T. Matsuda). The SP-shRNA BFP was created by replacing the GFP sequence by BFP2 sequence at the BspeI/BspeI sites. Luciferase constructs were obtained by PCR amplifying the 3’UTR regions from cDNA extracts and cloning them into a modified phRL-CMV vector (Promega), as previously described (Favereaux et al., 2011). For miRNA overexpression, a genomic sequence spanning 150 bp upstream and downstream of the miRNA sequence was PCR-amplified and cloned into a modified pcDNA3.1 with a U6 promoter (Invitrogen).

### Primary cultures and transfection

Primary rat hippocampal neurons were prepared from hippocampi of E18 Sprague-Dawley embryos (Janvier Labs, Saint Berthevin, France). Hippocampi were dissected out and processed for enzymatic dissociation for 15 min at 37°C in 0.05% trypsin-EDTA solution (Gibco) buffered with HEPES (Gibco) and containing 1% penicillin-streptomycin (100 mg.ml^-1^, Gibco). After washes, hippocampi were triturated with a serological pipette and around 450k cells were plated on glass coverslips (18 mm diameter, Marienfeld, 117 580) precoated with Poly-L-lysine hydrobromide (1 mg.ml^-1^, Sigma-Aldrich). Cells were placed into culture dishes containing Neurobasal™ medium (Gibco) supplemented with NeuroCult™ SM1 neuronal supplement (STEMCELL), L-glutamine (2 mM, PAA) and 3% horse serum (Invitrogen) at 37°C in 5% CO2. After 2-3 days in vitro (DIV), the culture medium was replaced with Neurobasal™ medium without horse serum and from DIV5-6 and every other 3-4 days, half of the medium was replaced by the same volume of BrainPhys™ medium (STEMCELL) supplemented with NeuroCult™ SM1 neuronal supplement (STEMCELL). Glial cells proliferation was inhibited by adding Ara-C (cytosine β-D-arabinofuranoside hydrochloride, 2.5 μM, Sigma-Aldrich) at DIV6-8. To induce HSP, half of the neuronal cultures were treated at DIV13 with 2 μM TTX (Tocris) for 48 h.

To test the effect of the knock-down of SP on AMPAR expression, neurons were cotransfected after 8 DIV with Homer1c-DsRed and either shRNA SP with GFP reporter or empty vector (pCDNA3-GFP) at a ratio of 1:4 and processed for immunocytochemistry at DIV14-16. For experiments using recombinant SEP-GluA2-containing AMPARs, neurons were co-transfected after DIV8 with SEP-GluA2 3’UTR-WT with SP-RFP at a ratio 1:1 and processed at 10 DIV. For miR-124 overexpression experiments, neurons were co-transfected after 10 DIV with Homer1c-GFP and either miR-124, miR-67 (miR control) or empty vector at a ratio of 1:3 using a lipofection protocol (Effecten, Qiagen) and processed at DIV15. To selectively prevent the interaction between miR-124 and SP-3’UTR, the rescue RFP-SP-3’UTR-WT / MUT constructs were cotransfected in cultured hippocampal neurons at DIV8 with Homer1c-BFP and SP-shRNA with GFP reporter before being processed at DIV14. A similar strategy was used to prevent the interaction between miR-124 and GluA2-3’UTR: dissociated hippocampal neurons were co-transfected at 8 DIV with Homer1c-DsRed and either SEP-GluA2 3’UTR-WT or SEP-GluA2 3’UTR-MUT at a ratio 1:3 and processed at DIV10. To inhibit glutamate synaptic release at individual inputs, neurons were transfected after DIV9-10 with EGFP-Synaptophysin:IRES:TetTx and processed at DIV13-14. To test the specificity of antibody against AMPA receptors, neurons were co-transfected at DIV9 with Homer1c-GFP and either HA-GluA1 or HA-GluA2 at a ratio 1:3 and processed at DIV10.

### Organotypic slices and whole-cell patch-clamp transfection

Organotypic hippocampal slice cultures were prepared from wild type mice (C57Bl6/J strain) as described (Stoppini et al., 1991). All animal experiments complied with all relevant ethical regulations. Animals were raised in our animal facility and they were handled and euthanized according to European ethical rules. Briefly, animals at postnatal day 5-8 were quickly decapitated and brains placed in ice-cold Gey’s balanced salt solution under sterile conditions. Hippocampi were dissected out and coronal slices (350 μm) were cut using a tissue chopper (McIlwain) and incubated at 35°C with horse serum-containing medium on Millicell culture inserts (CM, Millipore). The medium was replaced every 2-3 days. After 21 DIV, slices were transferred to an artificial cerebrospinal fluid (aCSF) containing (in mM): 130 NaCl, 2.5 KCl, 2.2 CaCl_2_, 1.5 MgCl_2_, 10 D-glucose, 10 Hepes (pH 7.35, osmolarity adjusted to 300 mOsm). Pairs of CA3 pyramidal cells were processed for whole-cell patch-clamp recordings to assess functional connectivity while infusing DNA constructs as previously described (Letellier et al., 2019). Recording pipettes were filled with a solution containing (in mM): 130 K-gluconate, 10 HEPES, 7 KCl, 0.05 EGTA, 2 Na2ATP, 2 MgATP, and 0.5 NaGTP (pH 7.30, osmolarity adjusted to 290 mOsm). During the recording, the presynaptic cell was infused with a plasmid encoding TetTx downstream of the synaptophysin-GFP and IRES sequences (100 ng.μl^-1^ in the recording solution) to label the silenced presynaptic terminals with GFP. Meanwhile, the postsynaptic cell was infused with plasmids encoding (1) shRNA-SP with a EBFP reporter (150ng.μl^-1^) allowing to knockdown endogenous SP while subsequently visualizing the cell morphology and (2) shRNA-resistant SP-RFP-3’UTR-WT or -MUT (150 ng.μl^-1^) to replace endogenous SP with a recombinant transcript containing or lacking the miR-124 binding region, respectively. After the recordings, the patch pipettes were gently retracted to facilitate membrane resealing. The slices were placed back in the incubator for 2 more days before being processed for confocal imaging.

### miRNA target prediction

Rat GluA2 mRNA sequence (encoded by the Gria2 gene) and SP mRNA sequence (encoded by the SYNPO gene) were PCR amplified and cloned from a brain cDNA library. MiRNA target prediction for the Gria2 and the SYNPO genes were performed with Targetscan algorithm (Agarwal et al., 2015).

### Total RNA isolation and qRT-PCR

For miRNA and mRNA quantifications, cultured neurons at 15 DIV were left untreated or treated for 48 h with 2 μM TTX and then processed for total RNA extraction using the kit Direct-zol™ RNA MicroPrep (Zymo Research) according to the manufacturer’s instructions. qRT-PCR was performed using miScript PCR System kit (Qiagen). Reverse transcription was achieved on 1 μg RNA with specific oligo-dT primers to generate cDNA library using this program: 60 min at 37°C followed by 5 min at 95°C and cooled down to 4°C. Quantitative PCR was carried out on a LightCycler LC480 (Roche) with primer pairs designed to span exon boundaries and to generate amplicons of ~100 bp. Primer sets for snRNA U6, GAPDH, Gria2, Synpo, miR-124, miR-92a, and miR-181a were tested by qRT-PCR and gel electrophoresis for the absence of primer-dimer artifacts and multiple products. Triplicate qRT-PCR reactions were done twice for each sample, using transcript-specific primers (600 nM) and cDNA (5 ng) in a final volume of 10 μl and using this program: 15 min at 95°C and 35 cycles of 15 sec at 95°C, 30 s at 55°C then 30 s at 70°C. The Ct value of each gene was normalized against that of snRNA U6 and GAPDH. The relative level of expression was calculated using the comparative (2-ΔΔCt) method (Schmittgen and Livak, 2008).

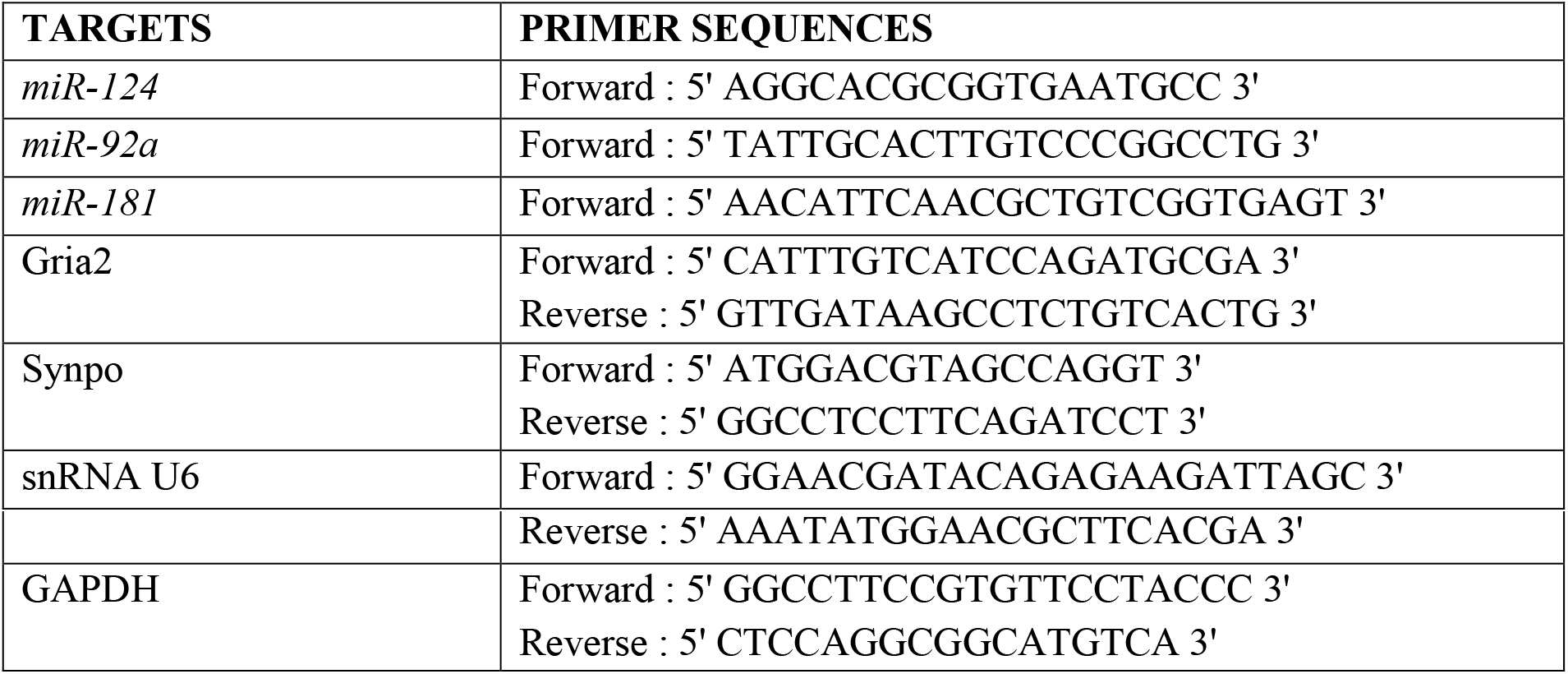

### HEK-293 cell culture and luciferase assay

HEK-293 cells were used to perform luciferase reporter assay. Cells were plated at 400k in a 25 cm^2^ flask (Dutcher) and cultured in Dulbecco’s modified Eagle’s medium supplemented with glutamax (3,87 mM), D-glucose (25 mM), penicillin (100 UI/mL) and streptomycin (100 mg/mL, Gibco). Cell cultures were maintained at 37°C in a humidified incubator with 5 % CO2. For luciferase reporter assay, HEK-293 cells were transfected with pmirGLO Dual-Luciferase miRNA Target Expression Vector (1 μg, Promega) containing a control reporter gene Renilla luciferase (hRluc) and either the 3’UTR of Gria2 or Synpo fused to the reporter gene Firefly luciferase (luc2). After 24 h, luciferase activity was measured using Dual-Luciferase Reporter Assay System kit (Promega) then normalized by the activity level of Renilla luciferase (hRluc).

### COS-7 cell culture and Western blotting

For Western blotting, COS-7 cells were plated at 120k cells per well in a 6-well plates (culture surface: 9,6 cm2 FALCON^®^, Dutscher) and cultured in Dulbecco’s modified Eagle’s medium supplemented with 10% SVF (Eurobio), 1% sodium pyruvate (Sigma-Aldrich), 1% glutamax (Gibco) and 1% penicillin-streptomycin (100 UI.ml-1 and 100 mg.ml-1, Gibco, Invitrogen), and maintained at 37°C in a humidified incubator with 5 % CO2. After 12-24 hours, cells were transfected with SP-RFP and either shRNA SP-GFP or empty vector (pCDNA3) using a lipofection method (X-tremeGENETM HP DNA Transfection Reagent, Roche) at a ratio 1:3. After 36-48 hours, cells were rinsed twice in ice cold PBS (ET330, Euromedex) and then scraped into 85 μL of RIPA buffer (50 mM Tris-HCl pH 7.5, 1 mM EDTA, 150 mM NaCl, 1% Triton-X100) containing protease inhibitor cocktail (Millipore). Homogenates were kept for 30 min on ice and then centrifuged at 8000xg for 15 min at 4°C to remove cell debris. The supernatant was recovered and the protein concentration was estimated using the Direct Detect^®^ Infrared Spectrophotometer (Millipore). 20 μg proteins were loaded on 4-15% Mini-PROTEAN^®^ TGX Precast Protein Gels Stain Free™ (Bio-Rad^®^) for separation (200 V, 400 mA, 40 min) and were subsequently transferred to nitrocellulose membranes for semi-dry immunoblotting (25 V, 1,3 A, 7 min, Bio-Rad^®^). Membranes were dried 5 min at 37°C and then incubated with 5% non-fat dried milk in Tris-buffered salineTween-20^®^ (TBST containing 28 mM Tris, 137 mM NaCl, 0.05% Tween-20^®^, pH 7.4) for 45 min at room temperature. Membranes were rinsed in TBST and cut to be incubated for 1 h at room temperature in 0.5% non-fat dried milk in TBST containing the appropriate primary antibody anti-SP (rabbit polyclonal, Synaptic Systems, 163 002, 1:1000), anti-Beta-actin (mouse monoclonal, Sigma-Aldrich, A5316, 1:5000) and anti-GFP (mouse polyclonal, Roche, 11867423001, 1:5000). After washing three times with TBST buffer, blots were incubated with horseradish peroxidase (HRP)-conjugated donkey anti-rabbit secondary antibody (Jackson Immunoresearch, 711-035-152, 1:5000) or HRP-conjugated donkey anti-mouse secondary antibody (Jackson Immunoresearch, 711-035-150, 1:10000) accordingly, in 0.5% non-fat dried milk in TBST for 1 h at room temperature. Target proteins were detected by chemiluminescence with Clarity™ (Beta-actin and GFP) and Clarity Max™ (SP) Western ECL Substrate kit (Bio-Rad^®^) on the Odyssey FC system (LI-COR). Average intensity values were calculated using Image Studio 5.2 software (LI-COR^®^). The intensity of SP signal was normalized by the intensity of beta-actin signal of the sample.

### Immunocytochemistry

For surface staining of AMPAR subunits, cultured hippocampal neurons were incubated live for 10 min at 37 °C with a mouse monoclonal antibody raised against the N-terminal domain of GluA2 that also recognizes GluA1 and GluA3 (Synaptic Systems, 182411 Clone 248B7) and diluted to 1:100 in the culture medium. For some experiments, neurons were incubated with a rabbit polyclonal antibody raised against the N-terminal domain of GluA1 (Agrobio, 2144 clone G02141) and diluted to 1:100 in the culture medium. Cells were subsequently fixed in paraformaldehyde (PFA) 4% with 4% sucrose for 10 min and permeabilized with 0,1% Triton X100 (Sigma-Aldrich) for 5 min. Endogenous SP, VGLUT1, MAP2 and/or PSD-95 were immunostained using primary antibodies diluted in PBS-BSA 1% for 30 min (SP: rabbit polyclonal, Synaptic System, 163 002, 1:600; VGLUT1: guinea pig polyclonal, Merck Millipore, AB5905, 1:5000; MAP2: rabbit polyclonal, Merck Millipore, AB5622, 1:1000; PSD-95: mouse monoclonal, Merck Millipore, MA1-046, clone 7E3-1B8, 1:500). After washes in PBS, neurons were incubated with PBS-BSA 1% (Sigma-Aldrich) for 30 min at room temperature and then stained with a secondary antibody conjugated to Alexa fluorophore diluted in PBS-BSA 1% at 1:800 for 30 min: anti-rabbit conjugated to either Alexa 350 (A11046, Thermo Fisher), Alexa 568 (A11011, Thermo Fisher) or Alexa 647 (A21244, Thermo Fisher); anti-guinea pig conjugated to Alexa 568 (A11075, Thermo Fisher); anti-mouse conjugated to either Alexa 488 (A11001, Thermo Fisher,), Alexa 568 (A11031, Thermo Fisher) or Alexa647 (A21236, Thermo Fisher). For surface staining of HA-GluA1/GluA2 or SEP-GluA2, neurons were incubated live for 10 min at 37 °C with anti-HA rat monoclonal antibody (1:100, 11867423001, Roche) or anti-GFP mouse monoclonal antibody (1:200, 11814460001, clones 7.1, 13.1, Roche), respectively. Cells were fixed and stained with anti-rat or anti-mouse secondary antibody conjugated to Alexa 568 (1:200, A11077, Thermo Fisher) or Alexa 647 (1:800, A21236, Thermo Fisher), respectively.

### *in situ* hybridization

For in situ hybridization of miR-124, double-digoxigenin locked nucleic acid probe (RNO-MIR-124-3P: CATTCACCGCGTGCCTTA, Tm: 84°C, 339111 YD00614870-BGC, miRCURY LNA™ miRNA Detection Probe, Qiagen) was used at a final concentration of 30 nM. Hybridization was performed according to the manufacturer’s instructions. Digoxigenin was revealed with a tyramide-based method (1:50, TSATM -Plus Fluorescein System, NEL741001KT, PerkinElmer). Pan neuronal staining was also processed (monoclonal mouse antibodies, 1:500, Neuro-ChromTM Pan Neuronal Marker Millipore MAB 2300) and revealed with a goat antimouse antibody conjugated to Alexa 568 (1:500, A11031, Thermo Fisher). Nuclei were stained with DAPI (DAPI Fluoromount-G^®^, SouthernBiotech).

### Epifluorescence microscopy and image analysis

Fluorescence imaging for immunocytochemistry and *in situ* hybridization on primary neuronal cultures was performed using an inverted microscope (Nikon Ti-E-Eclipse) equipped with a CMOS Prime 95B Scientific camera (Photometrics), an apochromatic (APO) x60/1.49 numerical aperture (NA) oil objective and filter sets allowing to image Alexa 350/DAPI (Excitation: FF01-379/34; Dichroic: FF-409Di03; Emission: FF-440/40; semROCK), GFP/Alexa 488 (Excitation: FF01-472/30; Dichroic: FF-495Di02; Emission: FF01-520/35; SemROCK), RFP/Alexa 568 (Excitation: FF01-543/22; Dichroic: FF-562Di02; Emission: FF01-593/40; SemROCK) and Alexa 647 (Excitation: FF02-628/40; Dichroic: FF-660Di02; Emission: FF01-692/40; SemROCK).

Image analysis was performed using Metamorph^®^ 7.8 software (Molecular Devices, Sunnyvale, USA). An intensity threshold was applied on Homer1c-GFP signal to define the neurite shape and then a binary segmentation was applied to define Homer1c-GFP clusters as synapses. For each synapse, the area and average intensity for both Homer1c-GFP and AMPAR signal were measured. Similarly, a binary segmentation was applied to the SP signal to define SP clusters. We then estimated the percentage of Homer1c-GFP clusters for which at least 20% of pixels was also positive for SP; these clusters were considered as SP+ synapses.

For experiments using TetTx expression in primary hippocampal cultures, VGLUT1 and MAP2 immunosignals were used to define glutamatergic presynaptic terminals and dendritic regions, respectively. A 14 x 14 pixels region surrounding VGLUT1 puncta was then defined. Within this region, integrated intensity for either AMPAR or SP signal was measured using Metamorph^®^ 7.8 software (Molecular Devices). 3 to 4 dendritic areas per neuron were analysed.

### FRAP experiment and analysis

FRAP experiments were performed on SEP-GluA2 signal for SP+ and SP-spines using an inverted microscope (Nikon Ti-E-Eclipse) equipped with an EMCCD camera (Evolve 512, Photometrics) driven by Metamorph^®^ software 7.10 (Molecular Devices), an APO x100/1.49 NA oil objective and filter sets to image SEP (Excitation: FF01-472/30; Dichroic: FF-495Di02; Emission: FF01-520/35) and RFP (Excitation: FF01-543/22; Dichroic: FF-562Di03; Emission: FF01-593/40). A laser bench comprising 491 and 561 lasers (100 mW each, Roper Scientific) was used to image or bleach SEP signal through a secondary optical fibre output connected to a device containing galvanometric scanning mirrors (ILAS, Roper Instrument). This device was driven by Metamorph^®^ software and allowed for precise spatial and temporal control of photobleaching in a user-defined targeted area. Switching between the two fibers for alternating between imaging and bleaching was performed in the millisecond range using a mirror. After acquiring a 30 s-baseline every 5 s, a rapid selective photobleaching of 6 to 10 spines (10 pixel-regions) was achieved using the 491 nm laser at higher level power (3 mW at the front of the objective) in less than 200 ms. Fluorescence recovery was then recorded immediately after the bleaching sequence during 750 s as follows: every 2.5 s for 50 s, every 5 s for 200 s and every 10 s for 500 s. Observational photobleaching was assessed by observing control nearby spines that were unbleached. Data were plotted as normalized fluorescence intensity vs time and fitted by a nonlinear regression (F = (1-IMf) (1-exp(-t/τ))) where F is SEP average fluorescence intensity, IMf is the immobile fraction and τ the time constant. Each value of average intensity for SEP signal was normalized by the average intensity of the baseline and by the average intensity of SEP signal for unbleached control spines. The time constant τ and immobile fraction IMf were calculated for 3 to 5 spines for each category of spine from several neurons.

### Confocal microscopy and image analysis

For visualization of putative synaptic contacts between CA3 transfected cells, organotypic slices were fixed in 4% PFA with 4% sucrose for > 4 h, washed in PBS, and subsequently mounted in Mowiol. Images were acquired on a commercial Leica TCS SP8 microscope (Leica Microsystems, Manheim, Germany) using a x 63/1.4 NA oil objective and a pinhole opened to 1 Airy unit. Images of 2 048 × 2 048 pixels, corresponding to a pixel size of 80-85 nm, were acquired at a scanning frequency of 400 Hz. The axial step size was set at 0.3 μm. Spine morphology was analyzed from 2D projections of confocal image stacks in ImageJ (NIH) using the custom-written plugin SpineJ as described (Letellier et al., 2019; Levet et al., 2020).

### mEPSC recording and analysis

Whole-cell voltage clamp recordings were made from 15-16 DIV hippocampal neurons at room temperature in a recording chamber continuously perfused at 2-3 ml.min^-1^ with extracellular aCSF solution (pH 7.3, osmolarity adjusted to 290 mOsm) and containing in mM (Sigma-Aldrich): 130 NaCl, 2.5 KCl, 2.2 CaCl_2_, 1.5 MgCl_2_, 10 D-glucose and 10 HEPES. aCSF was supplemented with bicuculline (20 μM, Tocris) and TTX (0.5 μM, Tocris). When stated, mEPSCs were recorded in aCSF supplemented with 10 μM *N*-[3-[4-(3-Aminopropylamino)butylamino]propyl]-2-naphthalen-1-ylacetamide trihydrochloride (NASPM, Tocris) to selectively block CP-AMPARs. Transfected neurons were identified with epifluoscence microscopy and differential interference contrast (DIC) illumination using an upright microscope (Nikon Eclipse FN1) equipped with Infinity 3s camera (Lumenera) driven by Metamorph^®^ software 7.13 (Molecular Devices) and with apochromatic x60/1.0 NA water objective. Patch pipettes with a resistance range of 5-6 MΩ were made from borosilicate glass capillaries (GC150T-10, Harvard Apparatus) using a vertical puller (PC-10, Narishige) and filled with an intracellular solution (pH 7.25, osmolarity adjusted to à 290 mOsm) containing in mM: 135 Cs-MeSO4, 8 CsCl, 10 HEPES, 4 MgATP, 0.3 NaGTP, 5 Qx-314-bromide and 0.3 EGTA. The electrodes were subsequently approached to target cells to achieved whole-cell patch-clamp using motorized micromanipulators (Scientifica). mEPSCs were recorded from neurons clamped at −70 mV using the multiclamp 700B amplifier (Molecular Devices), digitized at 10 kHz using a Digidata 1440A (Molecular Devices) and acquired using Clampex 10.3 (Molecular Devices). Access resistance (< 20 MΩ) was monitored every 2 min during the recording. Series resistance was left uncompensated. mEPSCs amplitude, frequency, 20-80% rise-time and decay-time constant were analyzed using MiniAnalysis software (v6.0, Synaptosoft).

### Statistical analysis

Graphs and statistical analyses were performed by using GraphPad Prism software. Outliers were identified with the ROUT method on raw experimental data. Data are presented as mean ± SEM of three or more experiments performed in independent preparations. Statistical differences were analyzed as indicated in figure captions. For normally distributed data (as determined by the Agostino-Pearson omnibus normality test), differences were tested using the two-tailed Student’s t test and one or two-way ANOVA test in case of three or more experimental groups. The Mann-Whitney and Kruskal-Wallis tests were used when criteria for normality were not met. Tukey and Dunn’s tests were used as multiple-comparison post hoc tests.

---

**Figure S1.**
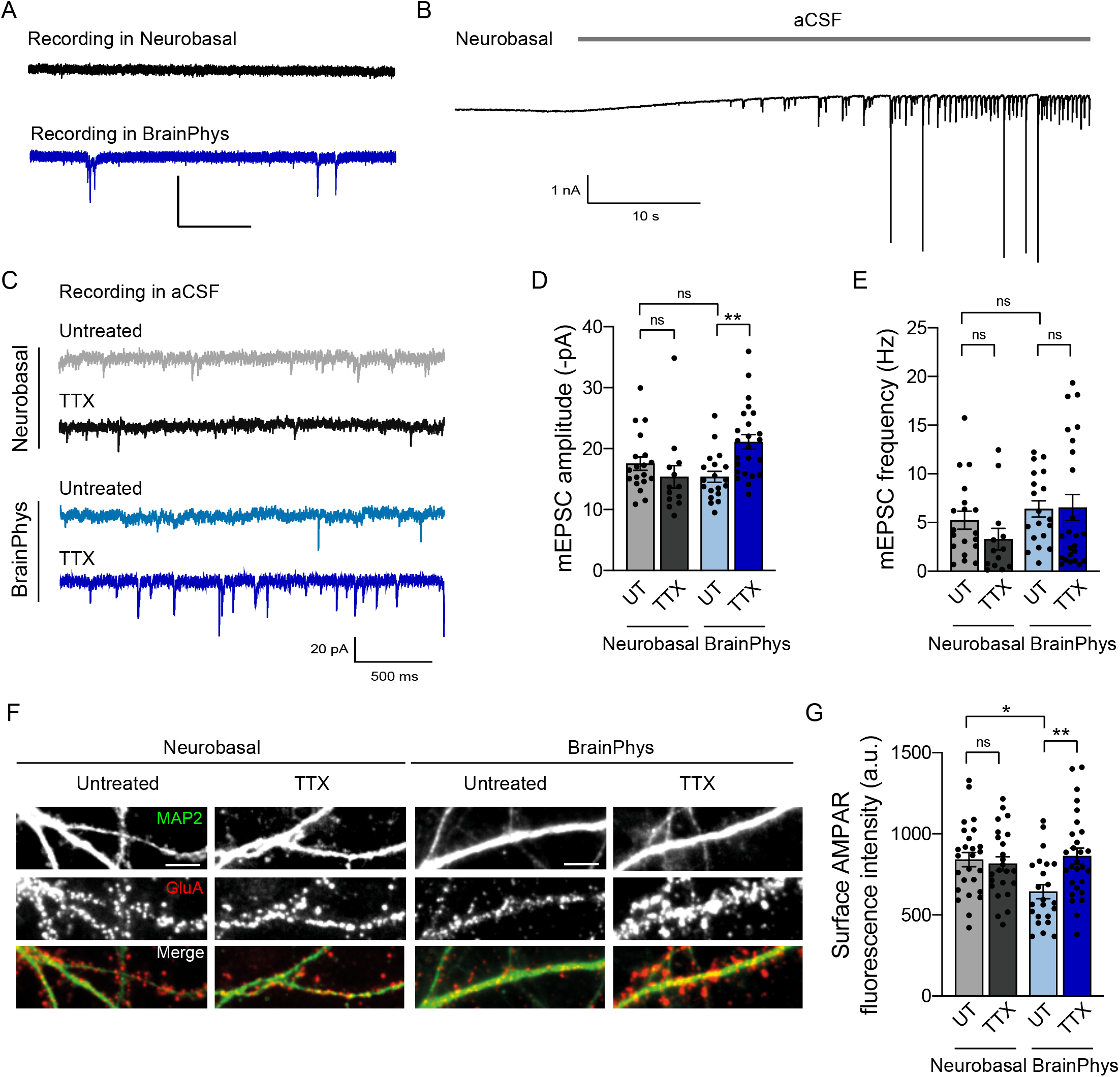
Culturing hippocampal neurons in Neurobasal-containing medium occludes HSP. **(A)** Representative traces of spontaneous synaptic currents recorded either in Neurobasal or BrainPhys-containing medium. **(B)** Recording of spontaneous activity from a neuron while washing-out Neurobasal-containing medium with articial cerebro-spinal fluid (aCSF). **(C)** Representative traces of AMPAR-mediated miniature currents (mEPSCs) recorded from neurons cultured in Neurobasal or BrainPhys and treated with TTX or not (untreated, UT). **(D,E)** AMPAR-mEPSC average amplitudes (D) and frequencies (E) for same condition as in (C) (Neurobasal: UT, n = 19, TTX, n = 13, BrainPhys: UT, n = 19, TTX, n = 24). n indicates the number of cells. **P < 0.01, ns, not significant, P > 0.05 by Kruskal-Wallis test followed by Dunn’s multiple comparison test. **(F)** Micrographs showing immunostaining for MAP2 (green) and surface endogenous AMPARs (red) for same conditions as in (C-E). Scale bar, 5 μm. **(G)** AMPAR synaptic fluorescence intensity for same condition as in (F) (Neurobasal: UT, n = 26, TTX, n = 24; BrainPhys: UT, n = 23, TTX, n = 28). n indicates the number of cells. **P < 0.01, *P < 0.05, ns, not significant, P > 0.05 by two-way ANOVA test followed by Tukey’s multiple comparison test. Data represent mean ± SEM.

**Figure S2.**
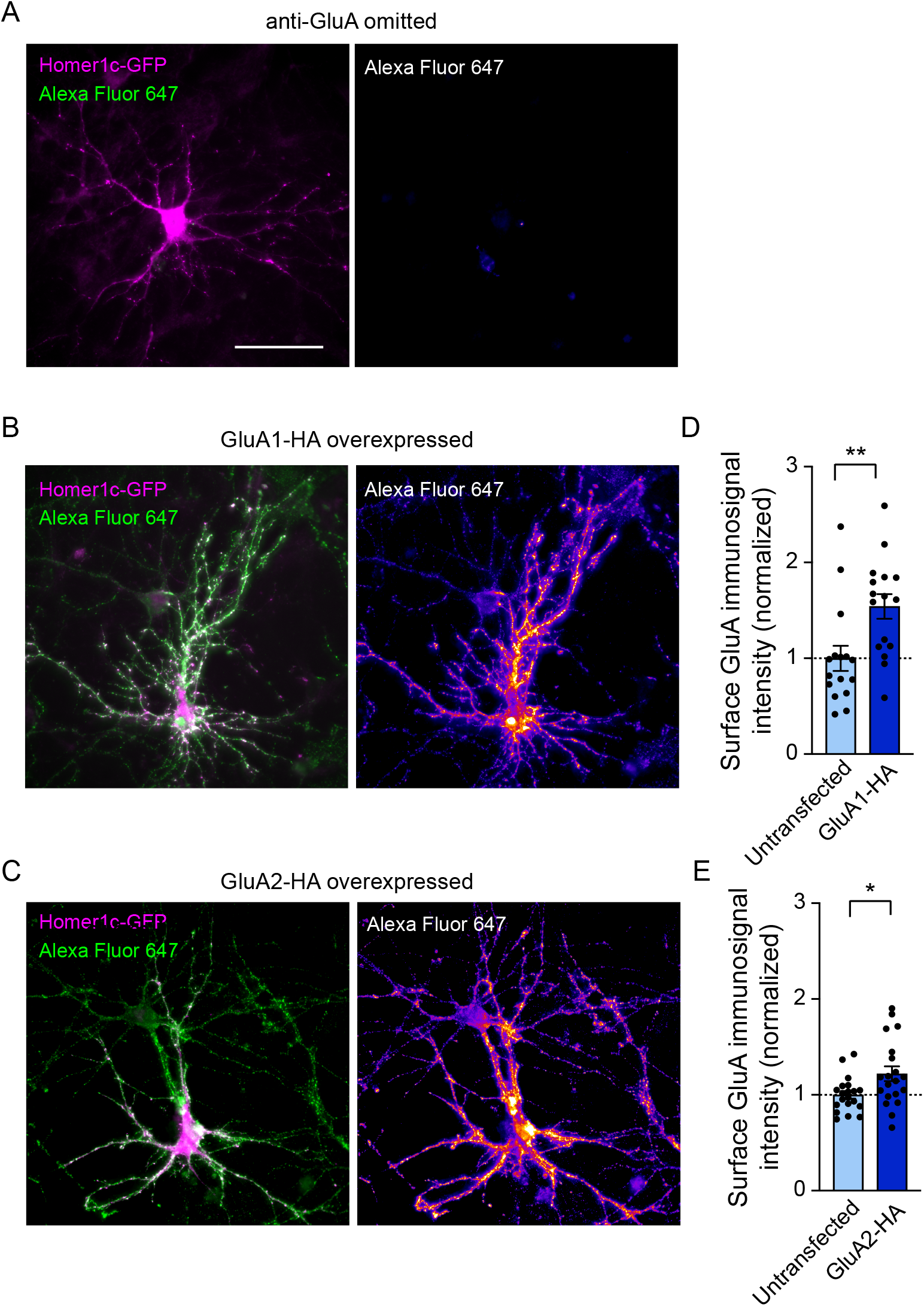
Specificity of AMPAR immunostaining in neurons. **(A)** Micrographs showing neurons transfected with Homer1c-GFP (magenta) when anti-GluA antibody was omitted before incubation with secondary antibody conjugated to AlexaFluor 647 (green). On the right panel, Alexa Fluor 647 signal has been coded with pseudo-colors. Scale bar, 60 μm. **(B,C)** Micrographs showing neurons transfected with Homer1c-GFP (magenta) and either GluA1-HA (B) or GluA2-HA (C), and immunostained for surface AMPARs with anti-GluA primary antibody and secondary antibody conjugated to Alexa Fluor 647 (green). On the right panel, Alexa Fluor 647 signal has been coded with pseudo-colors. Scale bar, 60 μm. **(D,E)** Quantification of surface GluA fluorescence intensity from untransfected vs transfected neurons expressing GluA1-HA (D) or GluA2-HA (E). GluA fluorescence intensity was normalized to untransfected condition (GluA1-HA: untransfected, n =16, transfected, n =16; GluA2-HA: untransfected, n = 20, transfected, n = 20). n represents the number of cells. GluA1-HA: **P = 0.0039 by Mann Whitney test; GluA2-HA: *P = 0.0160 by unpaired t test. Data represent mean ± SEM.

**Figure S3.**
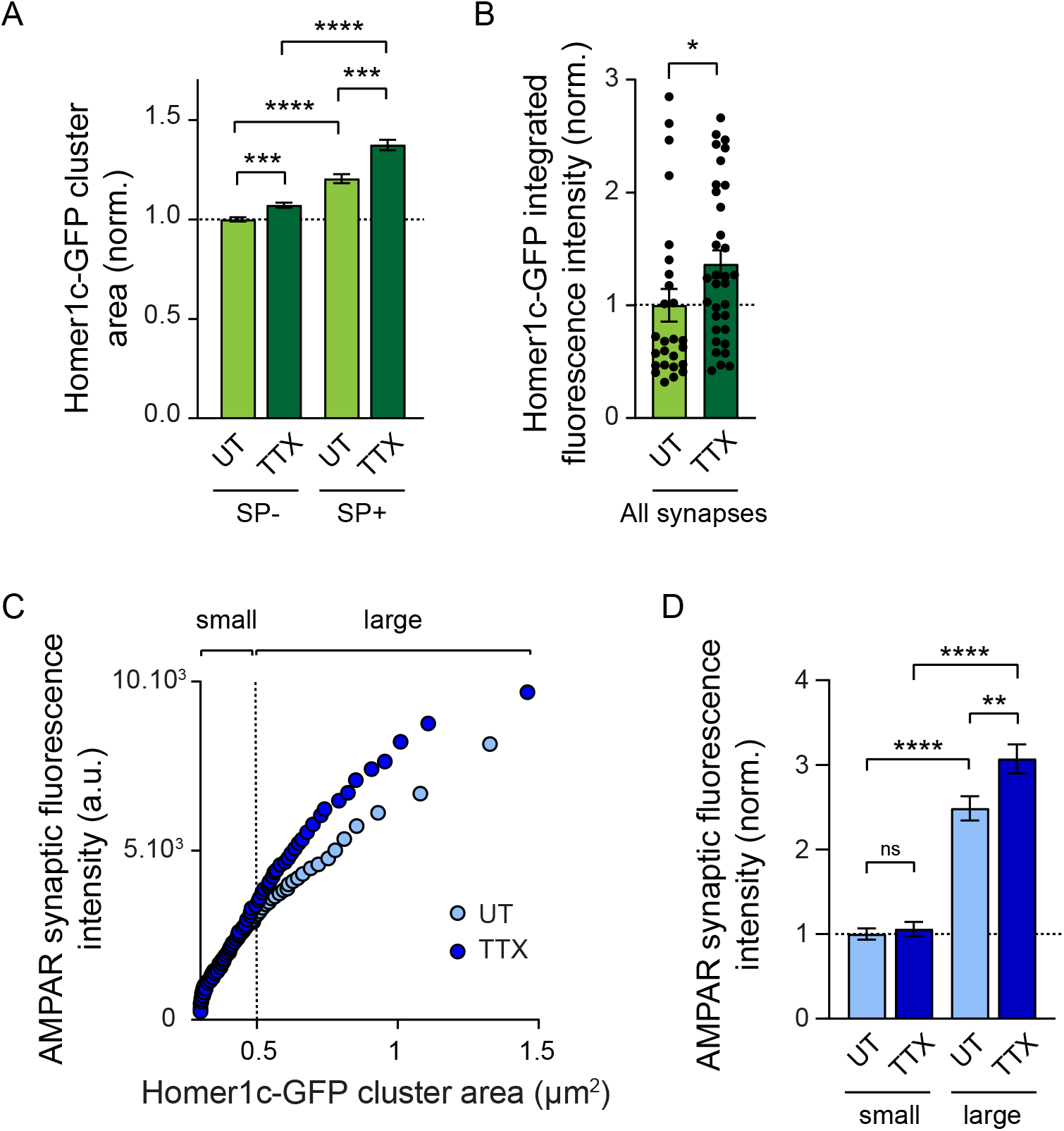
Synapse size correlates with SP expression and predicts synaptic AMPAR content depending on network activity. **(A)** Area of Homer1c-GFP clusters containing SP (SP+) or not (SP-) from neurons treated with TTX or left untreated (UT). Homer1c-GFP cluster area was normalized to untreated SP-synapse condition (SP-: UT, n = 1455, TTX, n = 1529; SP+: UT, n = 516, TTX, n = 724). n indicates the number of synapses. ***P < 0.001, ****P < 0.0001 by Kruskall-Wallis test followed by Dunn’s multiple comparison test. **(B)** Integrated fluorescence intensity of Homer1c-GFP clusters in UT or TTX-treated neurons, regardless of the expression of SP (all synapses). Homer1c-GFP cluster area was normalized to untreated condition. *P = 0.019 by Mann Whitney test. **(C)** Plots showing synaptic AMPAR fluorescence intensity vs synapse size for neurons treated with TTX (dark blue) or untreated (UT, light blue). The two curves were fitted using linear equations and the convergence of the traces to a common fit was tested using the extra sum of squares F test. The F test indicates that the traces are best fitted by two divergent linear models (P < 0.0001). Small and large synapses were defined according to the area of Homer1c-GFP clusters with a cut-off set at 0.5 μm^2^. **(D)** AMPAR synaptic fluorescence intensity at small vs large synapses in UT and TTX-treated neurons. AMPAR synaptic fluorescence intensity was normalized to small synapses from UT condition. **P < 0.01, ****P < 0.0001, ns, not significant, P > 0.05 by Kruskal-Wallis test followed by Dunn’s multiple comparison test. Data represent mean ± SEM.

**Figure S4:**
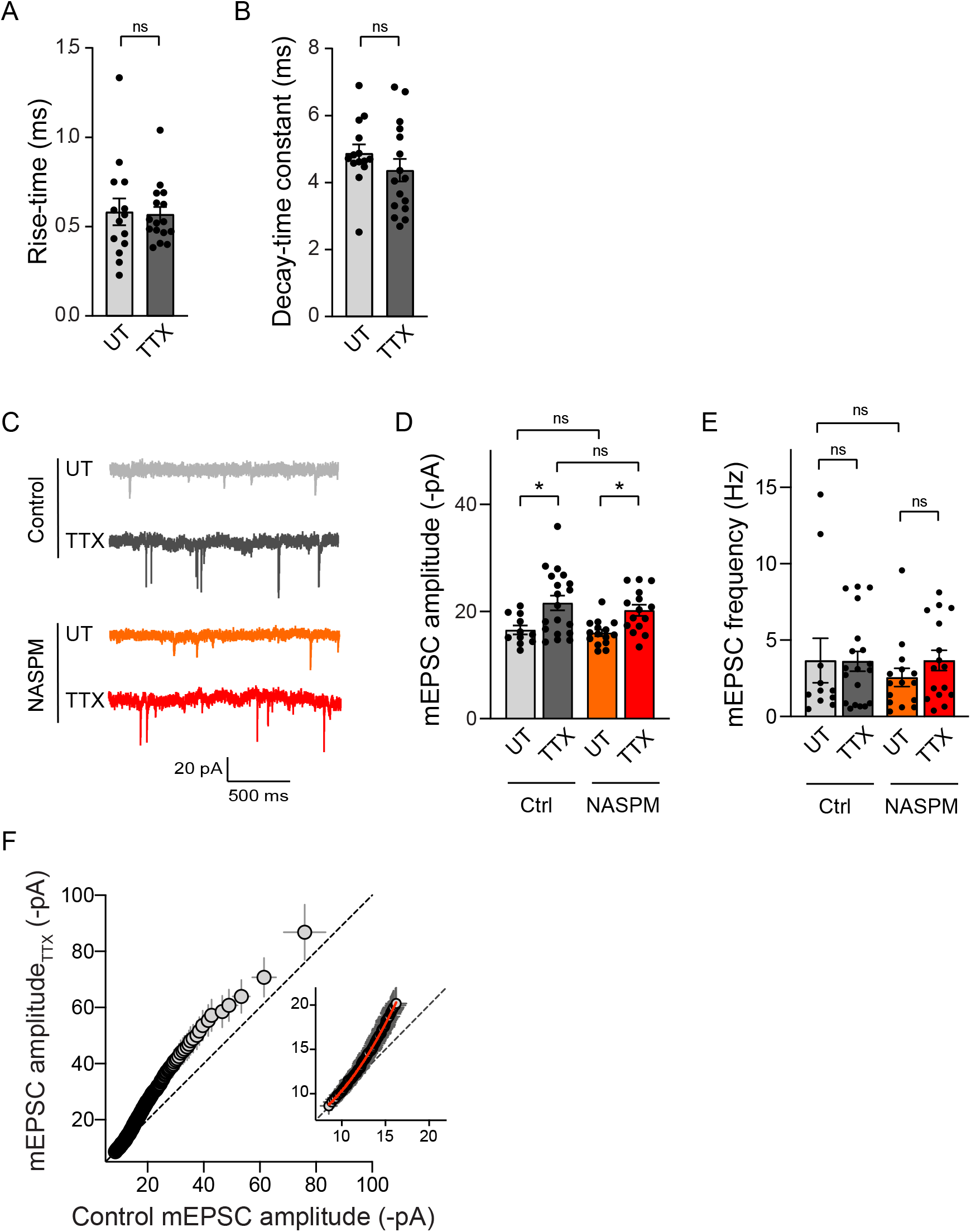
TTX-induced upscaling is not accompanied by change in mEPSC kinetics or sensitivity to NASPM. **(A,B)** Rise-time (A) and decay-time constant (B) for neurons treated with TTX or left untreated (UT) (UT: n = 14, TTX, n = 16). n indicates the number of cells. P > 0.05, ns, not significant by Mann Whitney test. **(C)** Representative traces of AMPAR-mediated mEPSCs from neurons treated with TTX, or untreated (UT) and recorded with or without (control, Ctrl) 10 μM NASPM. **(D,E)** Mean mEPSC amplitudes (D) and frequencies (E) for each condition (Ctrl: UT, n = 11, TTX, n = 19; NASPM: UT, n = 15, TTX, n = 15). n indicates the number of cells. *P < 0.05, ns, not significant, P > 0.05 by two-way ANOVA test followed by Tukey’s multi comparison test (D) or Kruskal-Wallis test (E). **(F)** Plot showing the rank-ordered AMPAR-mediated mEPSC amplitudes measured in TTX-treated neurons vs UT neurons (200 events). The rank-order plot was obtained by sorting from smallest to largest amplitude in untreated and TTX data and plotting them against each other. The extra sum of squares F test indicates that the first 100 events are better fitted with a second order polynomial quadratic curve (in red, TTX = 6.00 – 0.29 x UT + 0.07 x UT^2^; R^2^ = 0.99; ****P < 0.0001) than with a linear regression (not shown, TTX = −5.23 + 1.53 x UT). Data represents mean ± SEM.

**Figure S5.**
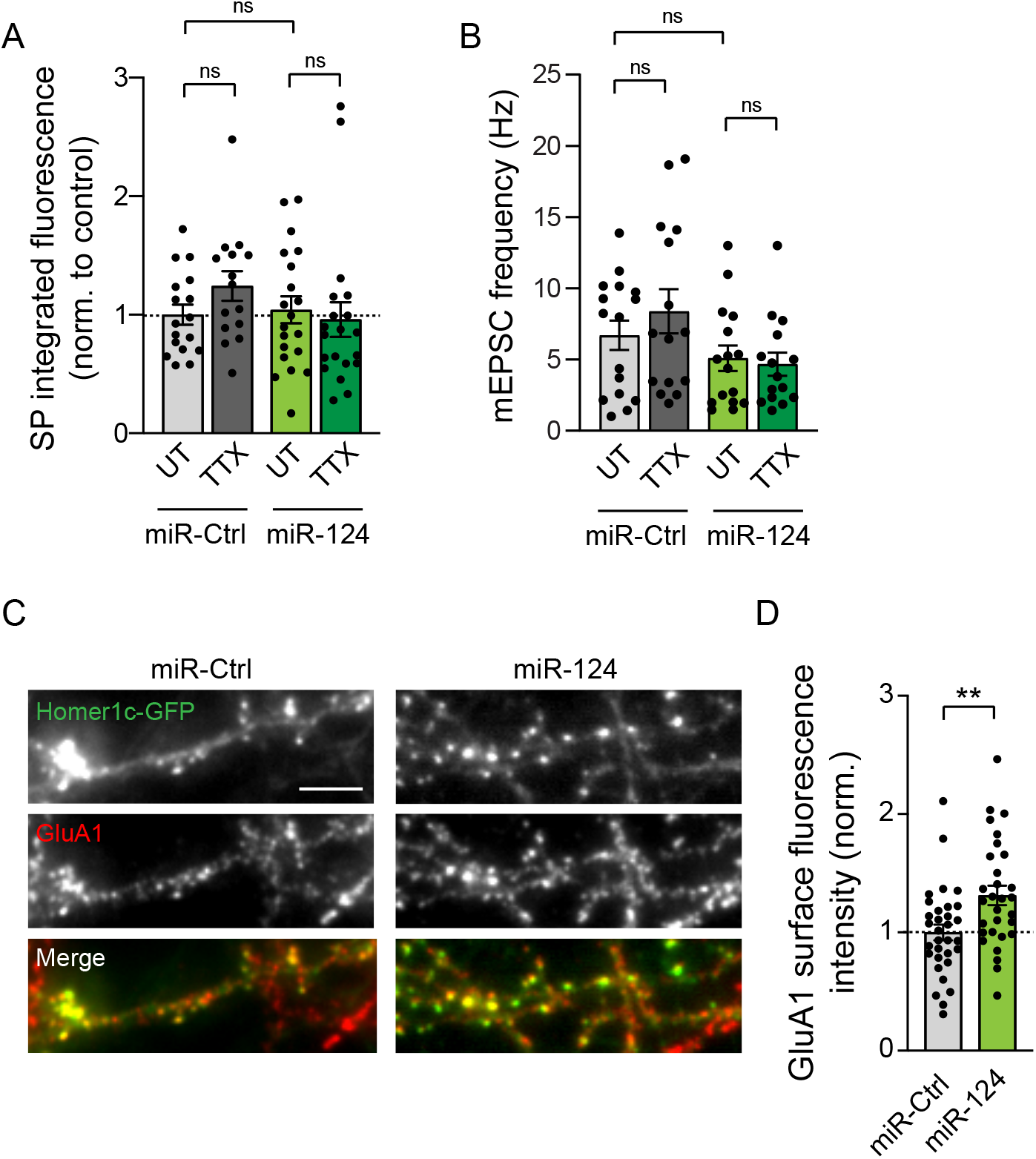
Effect of miR-124 overexpression on SP expression, mEPSCs frequency and GluA1 synaptic expression. **(A)** SP integrated fluorescence intensity in neurons expressing Homer1c-GFP and either miR-124 or control miR-67 (miR-Ctrl), and treated with TTX, or left untreated (UT). SP integrated fluorescence intensity was normalized to untreated miR-Ctrl condition (miR-Ctrl: UT, n =17, TTX, n = 15; miR-124: UT, n = 20, TTX, n = 20). n indicates the number of cells. ns, not significant, P > 0.05 by Kruskal-Wallis test. **(B)** mEPSCs frequency for same conditions as in (A). miR-Ctrl: UT, n = 16, TTX, n = 15, miR-124: UT, n = 16, TTX, n = 15). n indicates the number of cells. ns, not significant, P > 0.05 by Kruskal-Wallis test. **(C)** Micrographs showing neurons expressing Homer1c-GFP (green) and either miR-Ctrl or miR-124, and immunostained for surface GluA1 (red) in untreated neurons. Scale bar, 5 μm. **(D)** Surface GluA1 fluorescence intensity for same conditions as in (C). GluA1 surface intensity was normalized to miR-Ctrl condition (miR-Ctrl: n = 32; miR-124: n = 30). n indicates the number of cells. **P = 0.041, by two-tailed unpaired t test. Data represents mean ± SEM.

**Figure S6.**
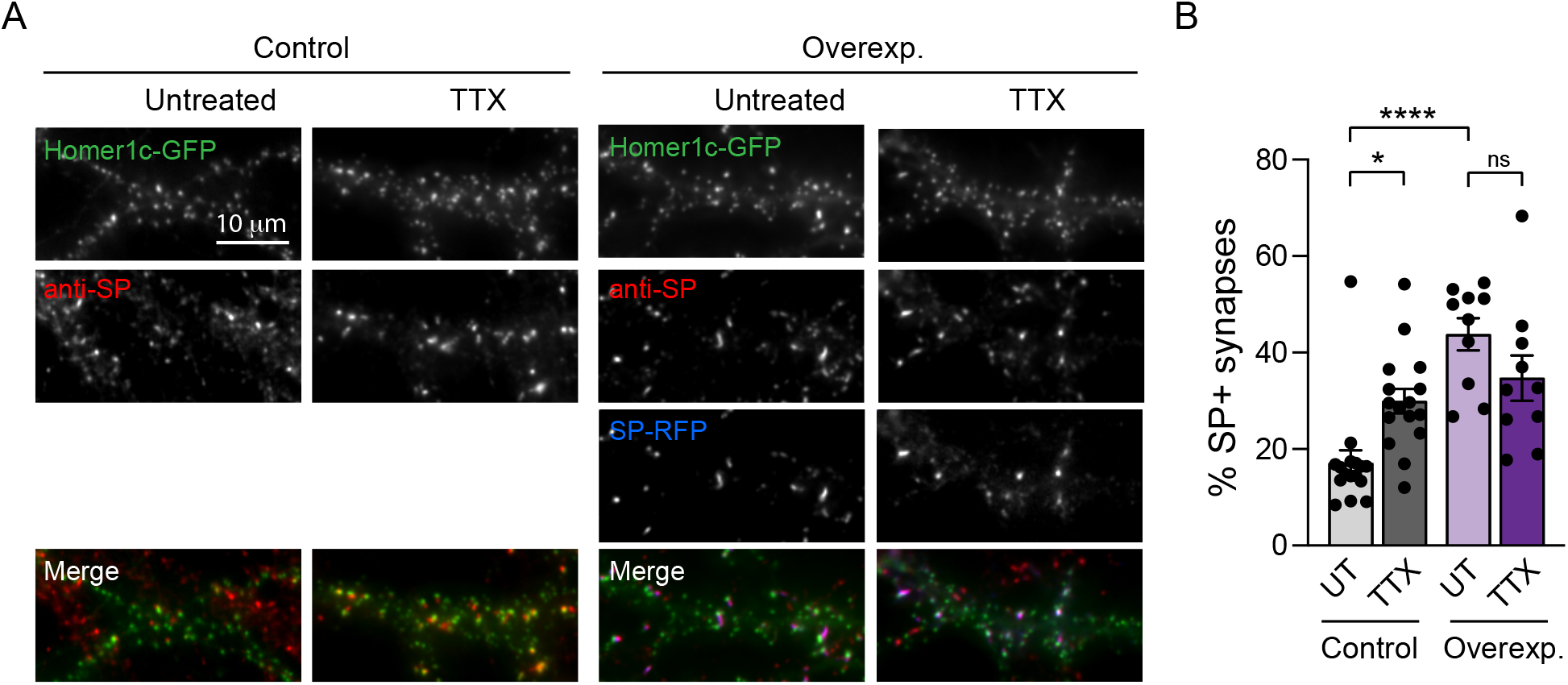
TTX-induced increase in the percentage of SP+ synapses is occluded by SP overexpression. **(A)** Micrographs showing transfected neurons with Homer1c-GFP alone (control, green) or with SP-RFP (overexp., blue), and treated with TTX, or left untreated (UT). SP (endogenous and exogenous) was immunostained using anti-SP antibody (red). Scale bar, 10 μm. **(B)** Percentage of SP+ synapses for each condition (Control: UT, n = 16, TTX, n = 16, Overexp.: UT, n = 10, TTX, n = 10). n indicates the number of the cells. *P < 0.05, ****P < 0.0001, ns, not significant, P > 0.05 by two-way ANOVA test followed by Tukey’s multiple comparison test. Data represent mean ± SEM.

**Figure S7.**
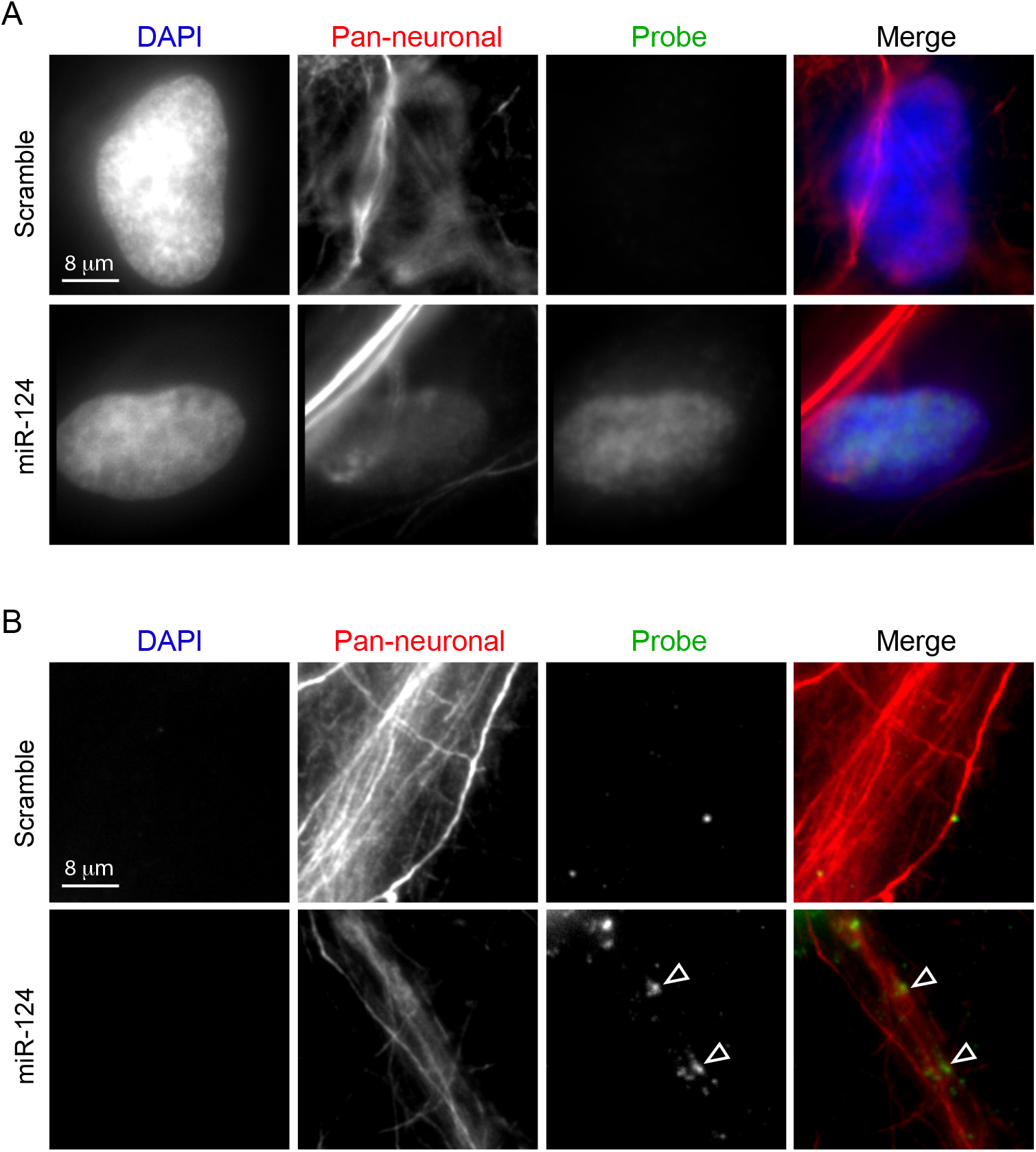
miR-124 is detected at both somatic and dendritic level in cultured hippocampal neurons. Micrographs showing fluorescent *in situ hybridization* of miR-124 or a control scramble probe (green) in soma **(A)** and dendrites **(B)** of neurons stained with DAPI (blue) and a pan-neuronal marker (red) to visualize neurites. Scale bar, 8 μm. Arrowheads indicate miR-124 puncta in neurites.

**Figure S8.**
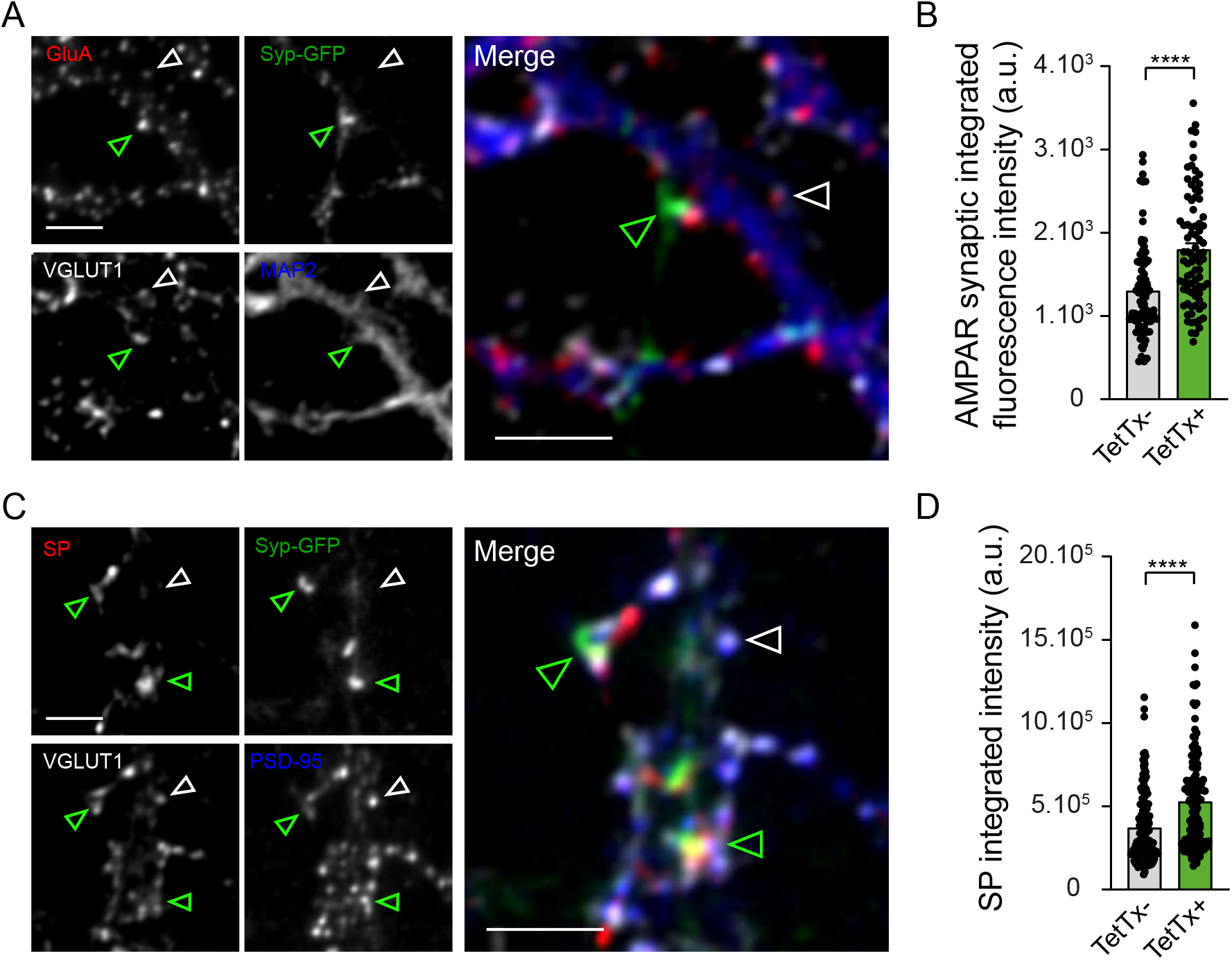
Synapse-autonomous homeostatic regulation of endogenous AMPARs and SP upon presynaptic silencing in cultured hippocampal neurons. **(A)** Micrographs showing dendrites from cultured hippocampal neurons immunostained for MAP2 (blue), VGLUT1 (grey) and AMPAR (red) and contacted by presynaptic terminals expressing Syp-GFP + TetTx (green). Arrowheads indicate GFP+ (green) and GFP-(white) terminals, immunopositive for VGLUT1. Scale bar, 5 μm. **(B)** AMPAR synaptic fluorescence intensity of clusters apposed to TetTx-vs TetTx+ terminals (n = 84 synapses for each conditions). ****P < 0.0001 by Mann Whitney test. **(C)** Micrographs showing dendrites from cultured hippocampal neurons immunostained for endogenous PSD-95 (blue), VGLUT1 (grey) and SP (red) and contacted by presynaptic terminals expressing Syp-GFP + TetTx (green). Arrowheads indicate GFP+ (green) and GFP- (white) terminals, immunopositive for VGLUT1. Scale bar, 5 μm. **(D)** SP integrated fluorescence intensity of clusters apposed to TetTx-vs TetTx+ terminals (TetTx-: n = 142; TetTx+: n = 143). n indicates the number of synapses. ****P < 0.0001 by Mann Whitney test. Data represent mean ± SEM.

**Figure S9.**
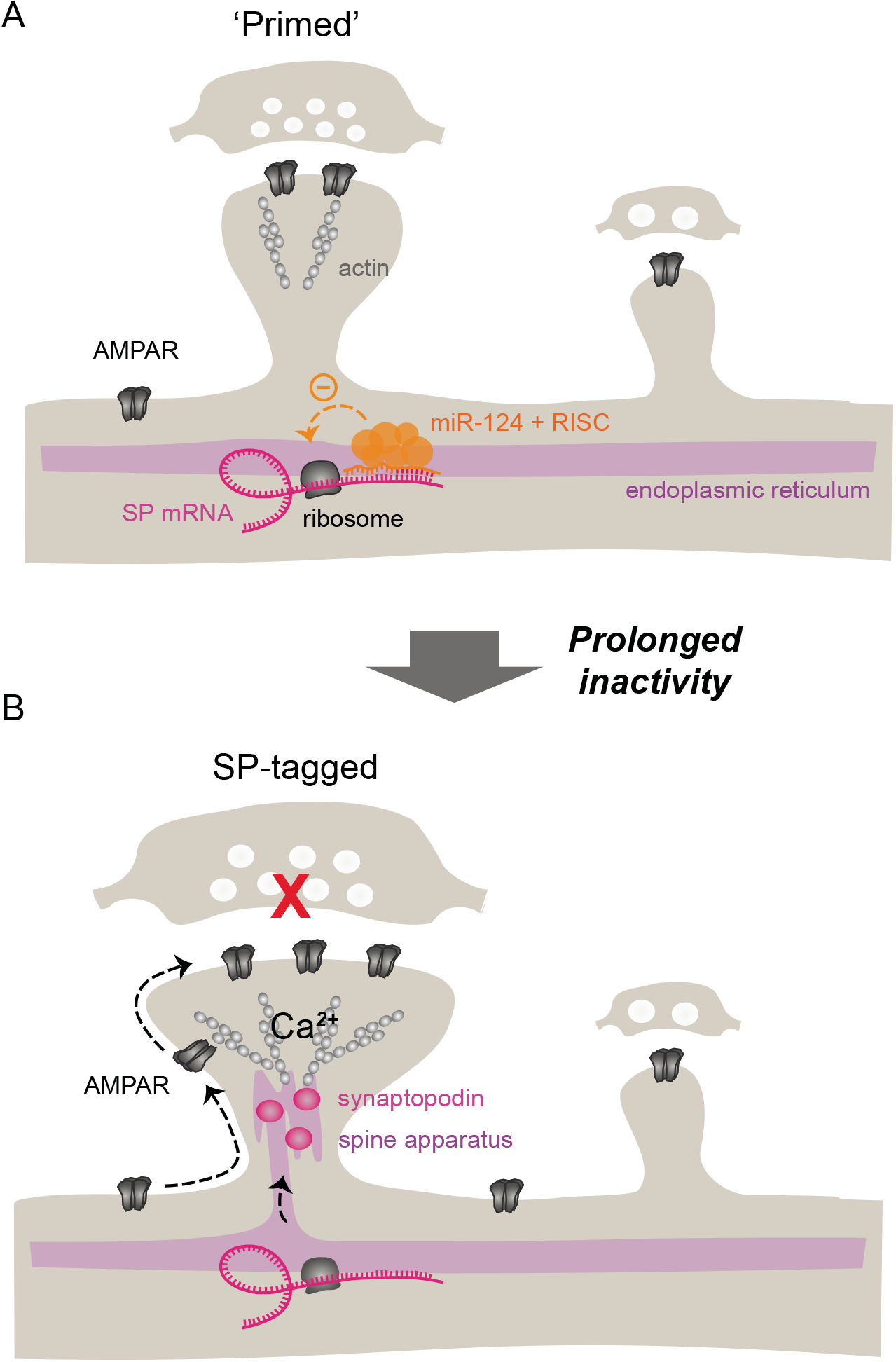
Working model. Under basal conditions, a subpopulation of synapses having SP transcripts bound to miR-124 at their proximity are ‘primed’ for HSP. Upon prolonged inactivity, SP translation by miR-124 is released at those synapses which get ‘tagged’ by SP. Synapse-autonomous SP expression promotes capture of surface diffusing AMPARs, spine growth and synaptic strengthening. RISC, RNA-induced silencing complex.

